# CORe Designer: a CRISPR design tool for proteome engineering

**DOI:** 10.64898/2026.06.12.731864

**Authors:** Akashaditya Das, Janessa Grech, Selina Hung, Jiali Wu, Celeste Callafe, Annabel Corrie, Kevin Yeung, Ione Goodwin, Daisy Yu, Sarah Hassan, Kimberley Reid, Marko Storch, Matthew Child

## Abstract

Precision proteome engineering requires tools that leverage CRISPR technologies, yet most current design platforms remain gene-centric. To bridge this gap, we introduce **CORe Designer**. Featuring an intuitive graphical user interface, **CORe Designer** enables researchers to design guide RNAs and homology-directed repair templates in a protein-centric manner. By automating these workflows, CORe Designer enables the design of targeted mutagenesis experiments for precision proteome engineering to be completed in minutes.

## Introduction

CRISPR-Cas9 is a powerful tool for precision genetic engineering in last decade. Advances in the field have made it possible to introduce genetic modifications at scale across diverse organisms, including larger programmed alterations and increasing numbers of simultaneous edits [1] [2] [3]. More recently, CRISPR-based therapies reached the clinic for the treatment of sickle cell disease and beta-thalassemia, while additional therapies for diseases including cancer and HIV are in clinical trials [4].

Despite CRISPR’s increasing utility, we believe the application of CRISPR Cas9 in precision proteome engineering is underutitilsed. We wanted to improve the workflow where a researcher makes targeted edits to a protein of interest to interrogate function. To accomplish this effectively, the researcher must first generate guide RNAs to target protein-coding sequences in a genome and initiate a Cas9 cleavage event. This is non-trivial when using currently available tools such as CHOPCHOP and CRISPETa [5] [6] as these popular tools are gene-centric requiring researchers to identify codon positions within genomes before they can be effectively used. While this is feasible for small numbers of guide RNA designs, this step becomes time consuming for larger studies. In the experimental design, a researcher must decide on how they want to edit the genome. Common strategies are to 1) rely on random edits introduced as a consequence of incorrect repair of the double-strand break following Cas9 cleavage; 2) use a Cas9 variant, such as one fused with an adenosine or cytosine base-editor for more predictable mutational outcome; 3) supply a donor template for homology directed repair in tandem with the Cas9 for fully customisable, programmed editing [7] [8] [9]. Of the three strategies, using donor templates for homology directed repair gives the most control over the edits that are made and this is what we decided to focus on. To simplify the design of programmed mutagenesis strategies using Cas9 with donor templates for homology directed repair, we built **CORe Designer**. This tool builds on two previous research outputs: **CRISPR-TAPE**, a guide RNA design tool that maps codons and identifies guides in a protein-centric manner, and our **CRISPR-based Oligo Recombineering (CORe)** method, which enables robust characterization of mutation tolerance in amino acid residues in proteins of interest [10] [11]. CORe Designer was built to run locally for quick generation of designs for protein mutagenesis and security over your sequences. It features a no-code graphical user interface for users to automate the creation of guide RNAs, donors for homology directed repair, and donor template specific primer pairs for enrichment of successful integration in downstream sequencing workflows (Figure 1a). We built CORe Designer to design mutagenesis experiments for a variety of organisms with minimal user inputs required. For guide RNA generation, the Uniprot ID, amino acid position and reference genome. For the generation of donor for homology directed repair adapted for our CORe methodology the user provides a codon usage table for their organism of choice. If the user’s organism has limited genomic information available, CORe designer can be run on smaller sections of genomic sequence input to generate guide RNA, donor templates and primer designs in the absence of a consensus genome (Figure 1b).

**Fig. 1:**
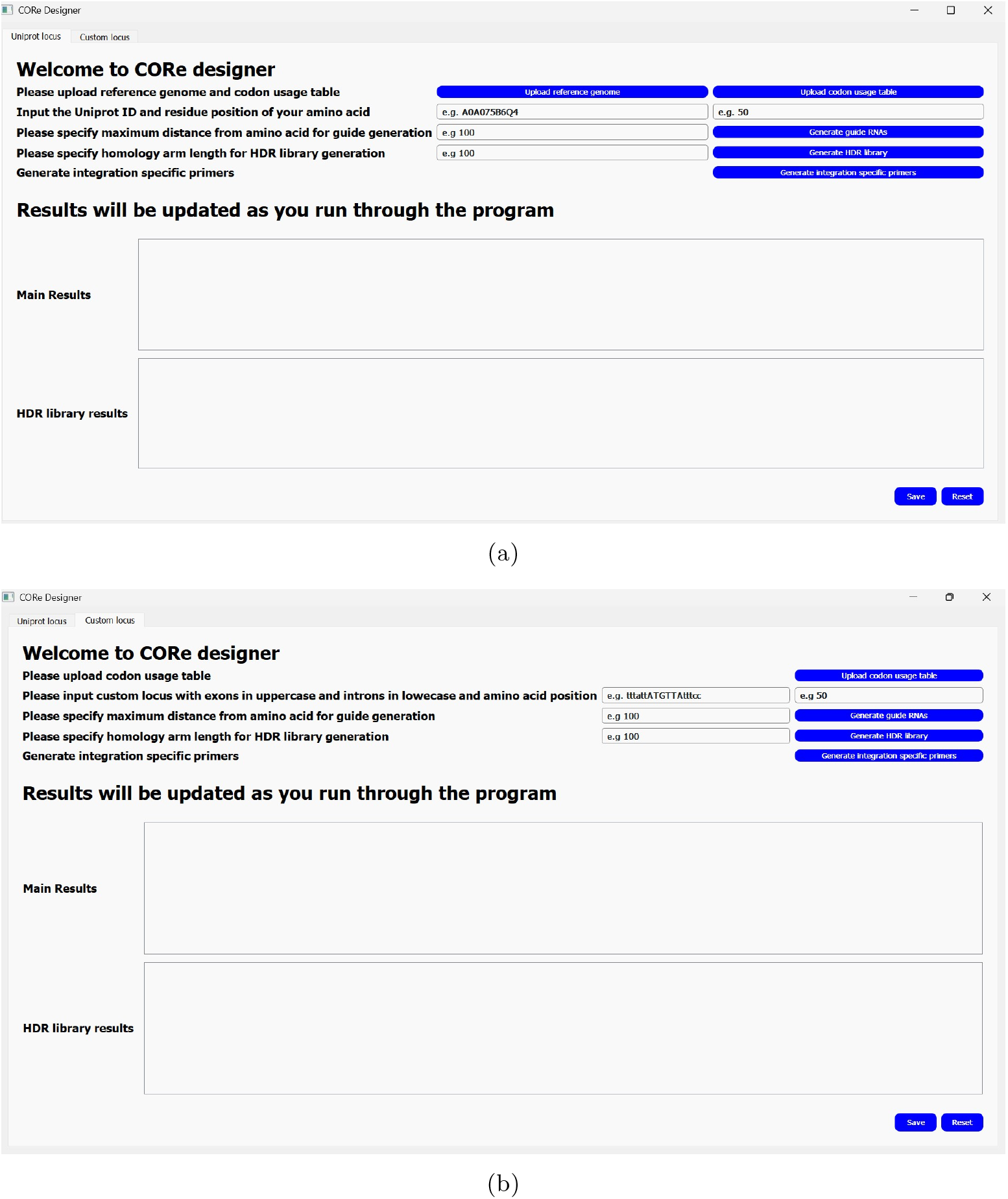
Screenshots of CORe Designer’s graphical user interface designed using PyQt5. a) Uniprot locus window that is used when genomic coordinates are linked to the protein of interest in Uniprot b) Custom locus window for where no genomic coordinates are available on Uniprot.

In this report, we pitch CORe Designer protein-centric tool for CRISPR mutagenesis experiments that sits alongside popular genome-centric tools [5] [6]. We provide all the code open-source so that readers use the tool as intended or extract snippets for their own workflows. We showcase CORe Designers performance and ease-of-use. As a demonstration of application, we then follow a CRISPR mutagenesis experiment from computational design using CORe to DNA synthesis of CORe Designer outputs to transfection of the DNA into cells and finally the sequence analysis of the mutated cells.

## Results and Discussion

### Key functions of CORe designer

In this section, we describe the CORe Designer processes (Figure 2a) and demonstrate their performance.

**Fig. 2:**
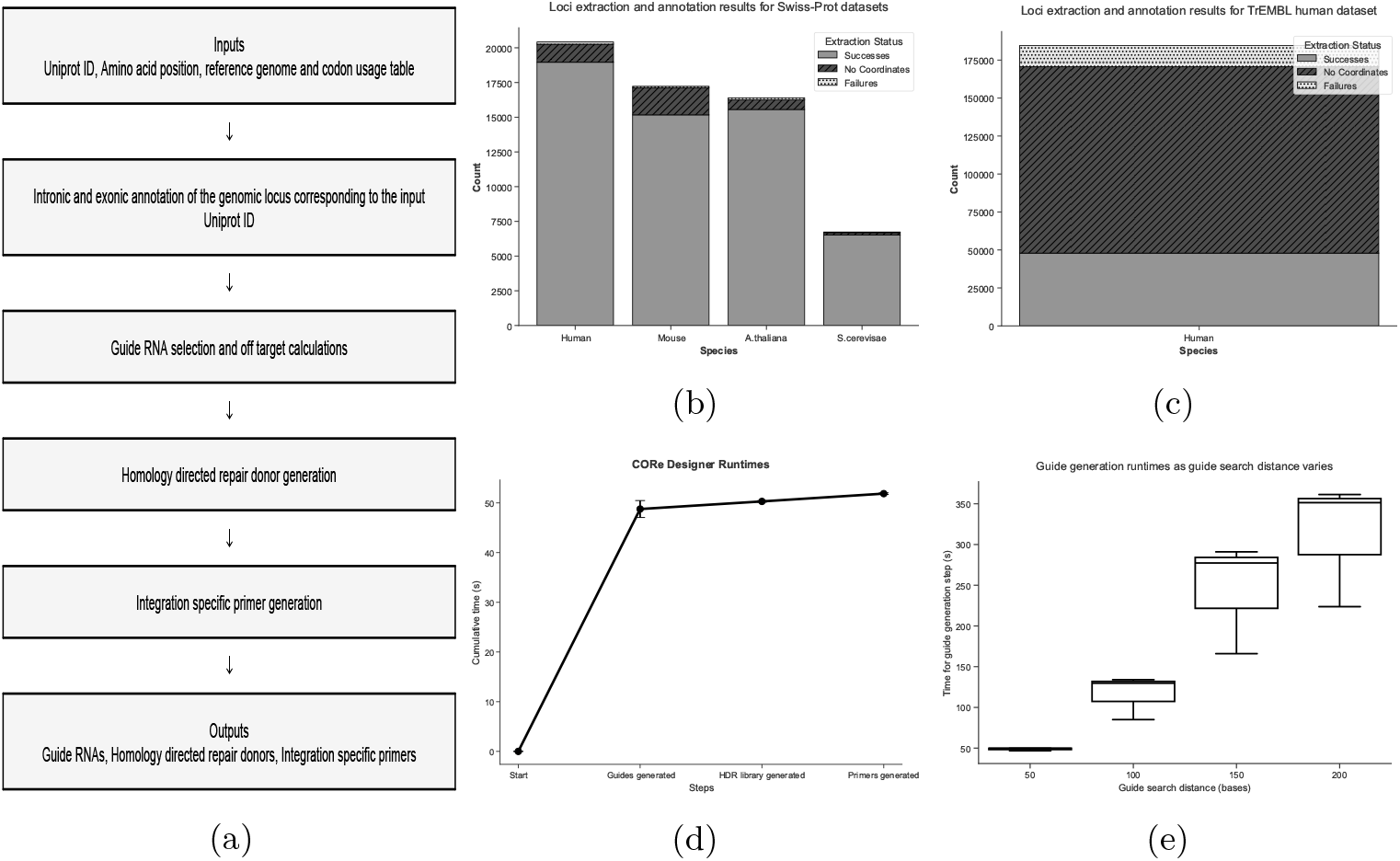
An overview of the features and the results of performance of CORe Designer. a) Schematic of the CORe Designer workflow. b) Bar chart detailing results of the automated loci identification and exon annotation on Swiss-Prot Uniprot entries. c) Bar chart detailing the results of the automated loci identification and exon annotation on trEMBL entries. d) Line plot detailing the time taken for each module of the CORe Designer workflow(n=3 and error bars represent standard deviation) e) Box plots detailing the time to find guides whilst varying the distance for guide RNA from the specified amino acid (n=3)

#### Automated genomic loci identification and exon annotation from Uniprot ID

The automated loci identification and exon annotation process uses the UniProt Proteins REST API coordinates function that uses a proteins UniProt ID as input [12]. This API output is retrieved in JSON format and converted into a Pandas DataFrame. A custom Python function that takes this DataFrame and the uses a reference genome to map it’s output to a DNA sequence corresponding to the protein Uniprot ID taken as input. Exonic sequences are defined in uppercase and intronic sequences in lower-case. To verify loci annotation, these exonic sequences are concatenated, translated into an amino acid sequence, and string matched against the protein sequence obtained by the API.

To test the effectiveness of this process, we ran these functions on all Swiss-Prot entries for four model organisms represented in UniProt: *Homo sapiens, Mus muscularis, Arabidopsis thaliana* and *Saccharomyces cerevisae*, testing a total of 60,781 sequences. We also ran the trEMBL entries of Homo sapiens to assess performance on a less stringently curated UniProt dataset, representing a total of 184,573 sequences. The results are presented in Figure 2b (Swiss-Prot) and Figure 2c (trEMBL). For the Swiss-Prot test cases with the four model organisms, 92.5% (56207/60781) passed the translation check. Of the UniProt IDs that failed, 6.8 % (4122/60781) were due to the absence of genomic coordinates associated with the entry, and the remaining 0.08% (452/60781) were due to the failure in the translation check of the extracted exons. We used *Homo sapiens* as trEMBL test case and found that a lower success rate of 26.0% (47903/184573) was achieved, with 66.5% of proteins with no associated genomic coordinates and 7.6% (13969/184573) failing in the translation check. trEMBL is a less stringently curated set of protein compared to Swiss-Prot so the lower success percentage was unsurprising. Translation check failures were surprising and we believe they are an edge case for the function or irregularities in the way that the information is inputted into the UniProt Database. We have attached lists of all the proteins checked and the code to generate this dataset in the supplementary folder 1.

To accommodate instances where extraction failure occurs, we have included an option in CORe Designer to input a custom locus with exons annotated in uppercase and introns in lowercase. Once the locus is supplied, the rest of the pipeline can be run with the caveat that genomic off-target counts cannot be calculated.

#### Guide RNA identification and selection

Guide RNA design was updated from the CRISPR-TAPE legacy code [11]. This process generates a codon mapping in the annotated locus where the position of each codon is identified. This mapping identifies the codon position for the user-inputted amino acid and maps that back onto the DNA locus for guide identification. The user defines a search window around the codon position in the loci where guide RNAs sequences are searched for. String matching is then identifies guide RNAs on both strands of DNA in the search window. For *Streptococcus pyrogenes* Cas9, gRNA sequences are defined as being 20 nucleotides immediately upstream of the NGG protospacer adjacent motif (PAM). Once guide RNA sequences are identified, features are calculated, including off target counts in the loci and genome. After the searches are complete, the user is prompted to select a pair of guide RNAs from those found. To benchmark the performance of this process, we calculated the run time for the workflow for example protein A0A075B6Q4 with residue position 50, a search window of 50 base pairs and donor for homology directed repair with homology arms of 100 base pairs. The search window and homology arm size were selected in line with previous CORe experiments where guide RNAs and donors for homology directed repair were generated manually. Run times were assessed and Figure 2d shows that guide generation was the computational bottleneck. Closer inspection suggested the majority of the compute time was for the search for genomic off targets. This is a common computational bottleneck for most guide RNA tools. Due to this, most tools are run on servers. As CORe Designer is run locally, we balanced the complexity of the off-target search to run in less than 5 minutes on a typical university-level laptop, we used a Dell Latitude 7420 Gen Intel(R) Core(TM) i7-1186G7 processor and 16GB of RAM. To assess the computational burden of this search, we increased the maximum guide search distance from 50 to 200 nucleotides, in increments of 50 nucleotides and recorded the time taken for this module in Figure 2e. This confirmed a linear dependence of compute-time vs search space and represented the range that we assume most users of CORe designer would use for CORe mutagenesis.

We recommend starting with a guide RNA search distance of 50 base pairs and increasing the maximum guide search distance only when no suitable guide RNA pairs are found.

#### HDR donor library preparation

After guide RNAs are selected, donor templates for homology directed can be automatically generated. The user is asked to define a homology arm length before a donor template is generated. The optimal length for these homology arms will be dependent on the size of the donor template being generated.

First, we determine whether the DNA sequence between the guide RNA pair is exonic, intronic or a combination of both. The distance between the base upstream of each guide RNA PAM and the codon coding for the amino acid of interest is calculated to identify the reference frame to ensure that no frame shifts are introduce during the recodonisation process. The recodonisation workflow converts each codon in the donor template to a synonymous codon with the closest codon frequency to the codon present in the wild type sequence. For the target amino acid, the codon of interest is replaced with the highest frequency codon for all 20 amino acids and a stop codon. This results in a library of 21 donor templates for each pair of selected guides. 5’ and 3’ homology arms are then appended to the recodonised donor sequences to create the donor library.

#### Integration specific primer design

In our CORe approach, regions flanking the target codon are synonymously recoded in the homology donor. This recoding generates new primer binding sites that are unique to genomes where the donor has been successfully integrated by homology directed repair. Priming off these recoded sections ensures that integration-specific amplicons can be selectively generated. We use a Python implementation of primer3 to generate primer sequences with these specifications, resulting in 600 base-pair amplicons with a melting temperature of 60°C.

### CORe improvements

In this section we detail improvements made to our CORe experimental workflow from Benns *et al* [10]. We use CORe designer to generate a mutagenesis experiement originally performed in Benns *et al* [10] and use our optimised CORe workflow to validate the design.

#### Donor libraries for homology directed repair can be stored in large oligonucleotide pools and extracted with PCR

The original application of CORe required ordering each donor template for each amino acid change as a separate piece of double-stranded synthetic DNA. These fragments represent each the different amino acid variants to be introduced by homology directed repair. Before transfection, these fragments needed to be manually mixed to the desired ratio. This method is time-consuming, introduces error between samples, and becomes cost-prohibitive as the number of targets required for a study increases. To enable affordable scale-up to multiple targets, we hypothesised that we could order large, complex oligo pools that encode for all donors for multiples and use PCR to selectively pull out the donor pool for the desired target. To achieve this, the donor library for a single target is flanked by a unique set of primer binding sites. For these primer binding sites we used the primer sites defined by Subramian*et al* which have previously been validated for orthogonality [13].

To test this strategy, we selected a target in *Toxoplasma gondii* that we have previously explored [10]; A cysteine residue at position 35 in RPL26 in *T*.*gondii* which we dub TgRPL26 C35. HDR donor pools were generated using CORe designer for nine other *T. gondii* targets alongside TgRPL C35 to give the pool diversity and better simulate a real use case for the amplification of an individual HDR donor subset from a larger pool.

The donor library specific to TgRPL26 C35 was selectively amplified from the full donor library pool. The identity of the amplified donor library was confirmed by Sanger sequencing (Figure 3c). As expected, Sanger sequencing revealed the expected sequence with ambiguous reads at the TgRPL26 C35 codon position, confirming sequence diversity at this codon position.

**Fig. 3:**
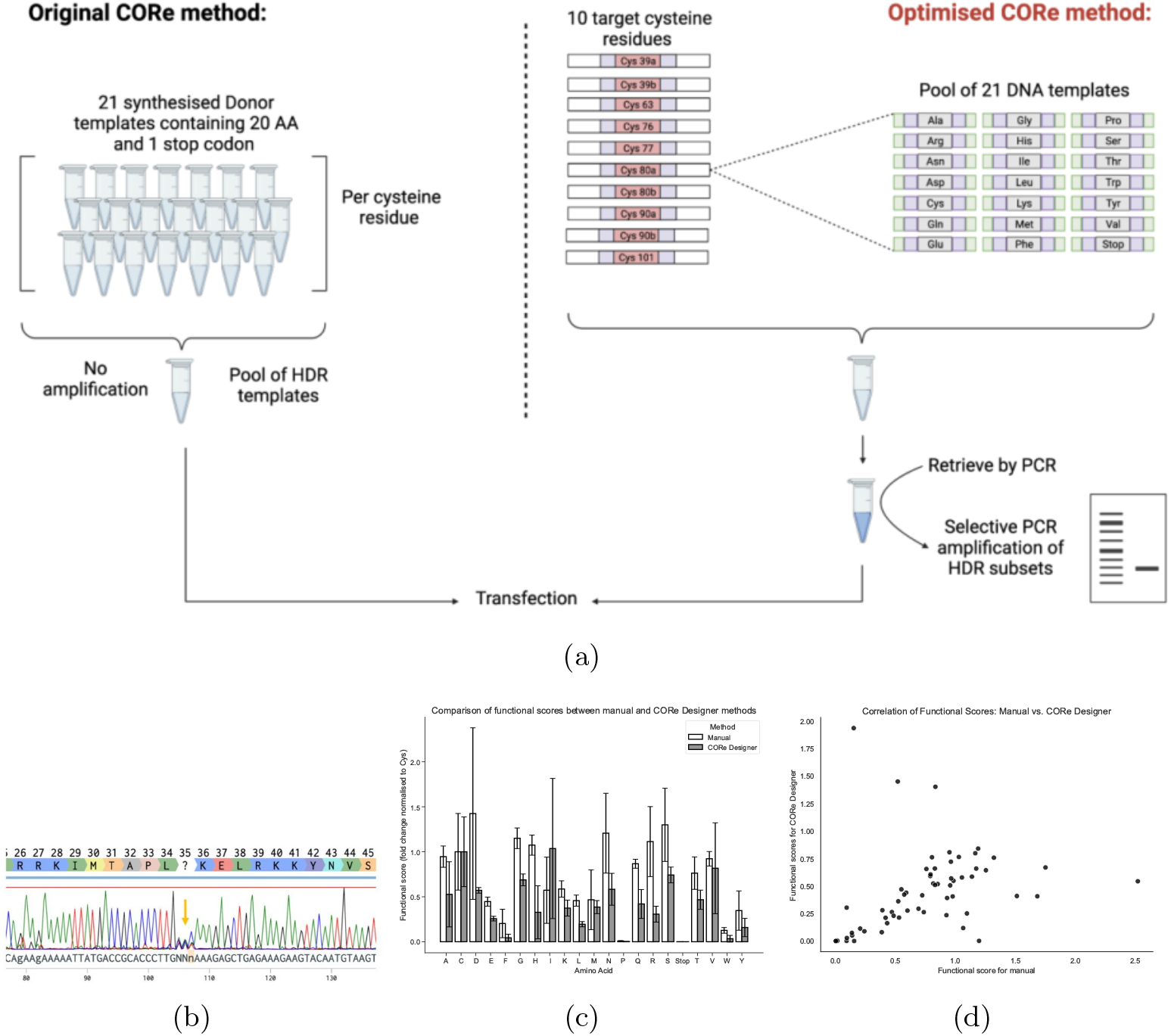
A walkthrough of how a CORe Designer design is taken from computational design to wet lab experimental outcome. a) Schematic showing the difference between the original CORe method and our optimised version. b) Sanger trace showing alignment of the recodonised region from an individual pool PCR from the larger, complex pool. c) Bar plot detailing the fitness scores (Fs) for each amino acid for the manual vs CORe designer designs. d) Scatter plot showing the Fs scores of the manual vs CORe Designer designs, for this plot replicate X manual pairs with replicate X CORe Designer where X = 1, 2, 3.

#### CORe Designer donors for HDR are comparable to manually designed donors for CRISPR-guided mutagenesis

The selectively amplified HDR donor subset for mutagenesis of TgRPL26 C35 was then co-transfected into *T. gondii* alongside the requisite CORe plasmid encoding Cas9 and guide RNAs targeting C35 in TgRPL26. DNA was extracted 1 day and 7 days after transfection for Illumina sequencing. We compared the mutagenesis effects of manually generated donor templates (as used in [10] and the donor designed by CORe designer. We then calculated Fs values as detailed in [10] and calculated the Pearson correlation coefficients of all the combinations of each manual and each automated replicate. Figure 3d shows a case where the functional score for Manual rep X plotted against CORe Designer rep X where X = 1,2,3. We calculated a mean coefficient of 0.65 across all combination of replicates. This shows that results from CORe Designer produces similar outputs to the manual design at a fraction of the labour cost.

## Conclusions

In this report we show that with only a user supplied Uniprot ID, amino acid position, reference genome and reference codon usage table CORe Designer is able to design CRISPR mutagenesis experiments that can be validated in the wet lab. This is achieved as a fraction of the labour cost that the designs would take to manually generate by a research scientist. We hope that CORe Designer to lower the access requirements for researchers around the world to run their own mutagenesis studies and are excited to see what you all will do with our tool.

## Methods

### Computational methods

#### CORe designer architecture

CORe designer is a graphical user interface application. The back-end is developed in a mixture of Python and C++ code, and the front-end is developed using PyQt5. The .exe was compiled using PyInstaller. We present our results from the standpoint of running CORe Designer using the Graphical User Interface provided as an executable file on Windows and Mac. A Readme is available in the Github alongside the .exe.

#### Performance testing

Performance testing was conducted Dell Latitude 7420 Gen Intel(R) Core(TM) i7-1186G7 processor and 16GB of RAM. Tests were run using CORe Designer app.py file in Windows with timing outputs written into the console. All code for runtime testing is provided in Supplementary file 1 and in the Github repository for CORe Designer.

### Experimental methods

#### DNA Synthesis

All DNA for guide RNA plasmids and HDR donor libraries were synthesised by Twist Biosciences. Primers for the CORe experiments were synthesised by Thermofisher Scientific.

#### Sub-library PCR

PCR reactions were performed using Q5 High-Fidelity DNA Polymerase (2X Master Mix)(NEB) in a 50µL reaction volume, containing 0.5µM of each forward and reverse primers and 2ng template DNA. Thermocycling conditions consisted of an initial denaturation at 98°C for 30s, followed by 35 cycles of 98°C for 10s, 65°C for 30s, and 72°C for 30s, with a final extension at 72°C for 120s. PCR products were assessed for fidelity with 2% agarose gel electrophoresis with SYBR Safe.

#### CORe mutagenesis

Transfections were carried out in 16-well Nucleocuvette strips using the Amaxa 4D-Nucleofector X-Unit (Lonza). Transfections were performed using the program ‘F1-115’. Transfections were carried out using freshly collected extracellular tachyzoites in P3 buffer (5 mM KCl, 15 mM MgCl2, 120 mM Na2HPO4/NaH2PO4 pH 7.2 and 50 mM d-mannitol). 7 *µ*g of pCORe-toxo (Genbank file provided in supplementary 2) was co-transfected with 1.05 *µ*g of the pooled mutational donors (representing 0.05*µ*g of each of the 21 mutational donors). Transfected parasites were expanded in HFF monolayers grown in 24-well plates and allowed to egress naturally 3 days after infection. Approximately 2 *×* 10^6^ of the egressed parasites were used to infect confluent HFF monolayers in six-well plates, and the remaining parasites (2 *×* 10^6^) were pelleted and frozen for genomic DNA extraction as the initial ‘pre’ mutant population. Parasites were allowed to egress naturally 5 days after infection and were similarly collected as the ‘post’ mutant population. Parasite genomic DNA from frozen cell pellets was extracted using the DNeasy Blood & Tissue Kit (Qiagen) for downstream NGS library preparation.

#### Illumina Sequencing

A 600bp fragment targeting the modified genomic locus was PCR amplified from parasite DNA. Primers were designed to include overhanging Illumina adapter sequences. The resulting amplicon was purified using AMPure XP magnetic beads (Beckman Coulter). Dual indices and sequencing adapters were ligated to the purified products using the Nextera XT Index Kit (Illumina). Indexed amplicons were then purified using AMPure XP beads, and quantified using the Qubit dsDNA HS/BR Assay Kits (Invitrogen). Indexed amplicons were pooled at equimolar concentration, and the size and purity of the resulting library assessed by gel electrophoresis. Pooled libraries were sequenced with Novogene using NovaSeq X Plus Series PE150 run.

#### Illumina Sequencing Data Analysis

Three biological replicates for the manually designed vs CORe designer mutagenesis experiments were assessed by Illumina sequencing. Fitness scores (Fs) were calculated using the same formula as in [10], where *F*_*s*_ = *f*_*post*_*/f*_*pre*_ where *f* = *reads*_*variant*_*/reads*_*allvariants*_. Pearson correlation coefficients were calculated on the Fs for each pair of replicates and the mean of these coefficients calculated. All code for data analysis is found in supplementary folder 1.

## Supporting information

Supplementary_folder_1

Supplementary_folder_2

## Declarations

- Funding: UKRI MRC Grant Ref: MR/Y031091/1
- Conflict of interest: No conflicts of interest
- Data availability: Illumina sequencing data generated and used in this project can be found in the NCBI SRA database under accession number PRJNA1477880.
- Code availability: All code used for this report can be found in the corresponding supplementary files. Code and executables for CORe Designer can be found at https://github.com/AkashD95/CORe-Designer
- Author contribution: A.D designed CORe Designer, analysed wet lab data and wrote the manuscript, J.G generated wet lab data and reviewed the manuscript, S.H and J.W created the first iteration of the app, C.C and A.C generated preliminary wet lab data. K.Y, I.G, D.Y, S.H, K.R tested CORe Designer and reviewed the manuscript. M.S and M.C provided grant funding, manuscript review and guided experimental design.

