## Supplementary_folder_1 for "CORe Designer: a CRISPR design tool for proteome engineering": Figure_1a.pdf

CORe Designer

Uniprot locus | Custom locus

### Welcome to CORe designer

Please upload reference genome and codon usage table — Upload reference genome — Upload codon usage table

Input the Uniprot ID and residue position of your amino acid — e.g. A0A075B6Q4 — e.g. 50

Please specify maximum distance from amino acid for guide generation — e.g 100 — Generate guide RNAs

Please specify homology arm length for HDR library generation — e.g 100 — Generate HDR library

Generate integration specific primers — Generate integration specific primers

### Results will be updated as you run through the program

**Main Results**

**HDR library results**

Save Reset
