## Supplementary_folder_1 for "CORe Designer: a CRISPR design tool for proteome engineering": 25_05_02_Human_Swiss_prot_batch_process_core_designs.ipynb

```
# %% [code]
#Import libraries and functions used in the notebook 
from Bio import SeqIO
from Bio.Seq import Seq
from Bio.SeqRecord import SeqRecord
from Bio.SeqFeature import SeqFeature, FeatureLocation
import os
import pandas as pd
from IPython.display import clear_output
import requests
import fnmatch 
import itertools
import primer3
from ahocorasick import Automaton

# List of folder paths to create
folders = [
    "batch_guides",
    "batch_selected_guides",
    "batch_hdr_templates",
    "batch_recodonised_locis",
    "batch_integration_specific_primers",
    "batch_genbank_files"
]

# Create each folder path
for folder in folders:
    os.makedirs(folder, exist_ok=True)  # exist_ok=True prevents errors if it exists
    print(f"Created: {folder}")

# %% [code]
#Utilty functions
def clean_loci(loci):
    """
    Cleans and processes loci of interest by removing introns, appending upstream and downstream sequences, and ensuring uppercase format.

    Arguments:
        loci (str) - input loci DNA sequence
        
    Returns:
        loci - cleaned loci sequence 
        loci_exon - exon sequence extracted from loci (assuming introns are coded as lower case)     
    """
    #Remove any leading or trailing blank spaces from loci
    loci = loci.strip()
    
    #Extract exons 
    loci_exon = ""
    for base in loci:
        if base.isupper():
            loci_exon += base

    return loci, loci_exon

def reverse_complement(dna_seq):
    """ 
    Return the reverse complement of a DNA sequence. 
    Arguments:
        dna_seq - DNA input sequence as a string
        
    Returns:
        reverse_complement - Reverse complement of DNA input sequence 
    
    """
    complement = {'a': 't', 'A':'T', 't': 'a', 'T':'A','c': 'g','C':'G', 'g': 'c', 'G':'C'}
    
    reverse_complement_seq = ''.join(complement.get(base, base) for base in reversed(dna_seq))
        
    return reverse_complement_seq

def translate(dna_seq):
    """
    Generates a protein sequence from a DNA sequence inputted as a string. It will only convert if the exon is in upper
    case

    Arguments:
        dna_seq(str) - input DNA sequence
        
    Returns:
        protein_sequence(str) - output protein sequence 
    """
    
    #Generate dictionary of codons for dna to protein translation
    
    codon_table = {
    'GCT': 'A', 'GCC': 'A', 'GCA': 'A', 'GCG': 'A',  # Alanine
    'TGT': 'C', 'TGC': 'C',                          # Cysteine
    'GAT': 'D', 'GAC': 'D',                          # Aspartic acid
    'GAA': 'E', 'GAG': 'E',                          # Glutamic acid
    'TTT': 'F', 'TTC': 'F',                          # Phenylalanine
    'GGT': 'G', 'GGC': 'G', 'GGA': 'G', 'GGG': 'G',  # Glycine
    'CAT': 'H', 'CAC': 'H',                          # Histidine
    'ATT': 'I', 'ATC': 'I', 'ATA': 'I',              # Isoleucine
    'AAA': 'K', 'AAG': 'K',                          # Lysine
    'TTA': 'L', 'TTG': 'L', 'CTT': 'L', 'CTC': 'L', 'CTA': 'L', 'CTG': 'L',  # Leucine
    'ATG': 'M',                                      # Methionine (Start codon)
    'AAT': 'N', 'AAC': 'N',                          # Asparagine
    'CCT': 'P', 'CCC': 'P', 'CCA': 'P', 'CCG': 'P',  # Proline
    'CAA': 'Q', 'CAG': 'Q',                          # Glutamine
    'CGT': 'R', 'CGC': 'R', 'CGA': 'R', 'CGG': 'R', 'AGA': 'R', 'AGG': 'R',  # Arginine
    'TCT': 'S', 'TCC': 'S', 'TCA': 'S', 'TCG': 'S', 'AGT': 'S', 'AGC': 'S',  # Serine
    'ACT': 'T', 'ACC': 'T', 'ACA': 'T', 'ACG': 'T',  # Threonine
    'GTT': 'V', 'GTC': 'V', 'GTA': 'V', 'GTG': 'V',  # Valine
    'TGG': 'W',                                      # Tryptophan
    'TAT': 'Y', 'TAC': 'Y',                          # Tyrosine
    'TAA': '*', 'TAG': '*', 'TGA': '*'               # Stop codons
    }
    
    # Initialize the protein sequence
    protein = ""

    # Iterate over the DNA sequence in steps of 3 (codon length)
    for i in range(0, len(dna_seq) - len(dna_seq) % 3, 3):
        
        # Define a codon as 3 bases
        codon = dna_seq[i:i + 3]
        # Translate the codon into an amino acid based on the dictionary
        if codon.isupper():
            protein += codon_table.get(codon, '?')  # '?' is a placeholder for unknown codons
        else:
            protein += 'x'

    return protein

def extract_genome_multifast_to_list_of_sequence_records(genome_multi_fasta):
    '''
    Takes in genomic multi fasta input and list of sequence records out 

    Args:
    genome_multi_fasta(fasta): multi fasta containing sequence records for a genome

    Returns:
    genome_sequence_records(list): list of sequences in biopython seqRecord format
    '''
    print('Starting genome extraction...')
    genome_sequence_records = []
    for record in SeqIO.parse(genome_multi_fasta, "fasta"):
        genome_sequence_records.append(record)
    print('Converted to list of sequence records')
    
    return genome_sequence_records

# %% [code]
#loci extraction
'''
This script will take in uniprot IDs and extract the genomic loci in format (introns in lowercase and exons in uppercase). This format is used by downstream processes to design guide RNAs,
homology directed repair templates and amplicon specific primers.
'''

def extract_genomic_information_from_uniprot_id(uniprot_id):
    '''
    Takes in uniprot ID as a string input and pings the Uniprot API to extract genomic coordinates of the protein and exons.
    Metadata such as name, taxID, protein sequence, genome assembly name,  ENSEMBL GeneID, ENSEMBL Transcript ID and ENSEMBL Translations IDs is included alongside the extracted coordinates.  

    Args:
    uniprot_id (str): uniprot ID

    Returns:
    genomic_information (pd.DataFrame): DataFrame containing genomic coordinates of the protein of interest alongside exon positions and metadata 
    '''
    extraction_failure = False
    genomic_information = pd.DataFrame()
    try:
        print(f'Searching for UniProt ID: {uniprot_id}')
        requestURL_protein = f"https://www.ebi.ac.uk/proteins/api/coordinates/{uniprot_id}"
        response_protein = requests.get(requestURL_protein, headers={"Accept": "application/json"})
        
        # Check if the request was successful
        response_protein.raise_for_status()
        
        # Load JSON response
        response_protein = response_protein.json()
        
        # Check if response is not empty
        if response_protein:
            response_protein_normalise = pd.json_normalize(
                response_protein, 
                record_path=['gnCoordinate', 'genomicLocation', 'exon'], 
                meta=['accession', 'name', 'taxid', 'sequence', 
                      ['gnCoordinate', 'genomicLocation', 'chromosome'], 
                      ['gnCoordinate', 'genomicLocation', 'start'], 
                      ['gnCoordinate', 'genomicLocation', 'end'], 
                      ['gnCoordinate', 'genomicLocation', 'reverseStrand'], 
                      ['gnCoordinate', 'genomicLocation', 'nucleotideId'], 
                      ['gnCoordinate', 'genomicLocation', 'assemblyName'], 
                      ['gnCoordinate', 'ensemblGeneId'], 
                      ['gnCoordinate', 'ensemblTranscriptId'], 
                      ['gnCoordinate', 'ensemblTranslationId']],
                record_prefix='exon_'
            )

            # Group and aggregate exon information
            response_protein_normalise = response_protein_normalise.groupby([
                'accession', 'name', 'taxid', 'sequence', 
                'gnCoordinate.genomicLocation.chromosome', 
                'gnCoordinate.genomicLocation.start', 
                'gnCoordinate.genomicLocation.end', 
                'gnCoordinate.genomicLocation.reverseStrand', 
                'gnCoordinate.genomicLocation.nucleotideId', 
                'gnCoordinate.genomicLocation.assemblyName', 
                'gnCoordinate.ensemblGeneId', 
                'gnCoordinate.ensemblTranscriptId', 
                'gnCoordinate.ensemblTranslationId'
            ]).agg({
                'exon_id': lambda x: ','.join(map(str, x)),
                'exon_proteinLocation.begin.position': lambda x: ','.join(map(str, x)),                    
                'exon_proteinLocation.end.position': lambda x: ','.join(map(str, x)),
                'exon_genomeLocation.begin.position': lambda x: ','.join(map(str, x)),                    
                'exon_genomeLocation.end.position': lambda x: ','.join(map(str, x))
            }).reset_index()

            # Concatenate to the main DataFrame
            genomic_information = pd.concat([genomic_information, response_protein_normalise], ignore_index=True)
        else:
            print(f"No data found for UniProt ID: {uniprot_id}")
            
    except Exception as e:
        print(f"An error occurred: {e}")
        extraction_failure = True
        print(f'Failed extractions:{extraction_failure}')

    return genomic_information, extraction_failure

def recursive_translate(genomic_loci_exons, genomic_information_loci_extracted):
    print('recursive translation...')
    # Initial check if "ATG" exists
    index = genomic_loci_exons.find("ATG")
    
    # If "ATG" is not found, return a fail
    if index == -1:
        genomic_information_loci_extracted = genomic_information_loci_extracted.assign(translation_check='Fail')  
        print("ATG not found in the sequence")
        print(genomic_information_loci_extracted) 
        return genomic_information_loci_extracted

    # List to keep track of all ATG positions
    atg_positions = []
    while index != -1:
        atg_positions.append(index)
        # Search for the next "ATG" after the current index
        index = genomic_loci_exons.find("ATG", index + 1)

    # Now, iterate through each ATG position and try translating
    for atg_index in atg_positions:
        atg_start_sequence = genomic_loci_exons[atg_index:]
        #print(f"Translating from ATG position {atg_index} /coding DNA length({len(atg_start_sequence)})...")
        #print(atg_start_sequence)
        translation = translate(atg_start_sequence)
        #print(f"Translation length ({len(translation)}): {translation}")
        # Check if translation matches the expected sequence
        if translation == genomic_information_loci_extracted['sequence'].iloc[0]:
            genomic_information_loci_extracted = genomic_information_loci_extracted.assign(translation_check='Pass')   
            print("Translation matched. Marking as 'Pass'.")
            print(genomic_information_loci_extracted)
            #genomic_information_loci_extracted['loci_exons'] = atg_start_sequence
            return genomic_information_loci_extracted

        else:
            continue
            #print('Translation match failed. Checking next index position.')
    # If no translation matches after checking all positions, mark as 'Fail'
    genomic_information_loci_extracted = genomic_information_loci_extracted.assign(translation_check='Fail')   
    print("No matching translation found. Marking as 'Fail'.")
    print(genomic_information_loci_extracted)
    return genomic_information_loci_extracted

def extract_genomic_loci_from_genomic_information(genomic_information, genome_sequence_records, UTR_length=1000):
    '''                  
    Takes the output from the Uniprot API from extract_genomic_information_from_uniprot_id function and extracts the genomic loci using a given reference genome. It appends the genomic loci, 
    genomic loci exons and performs a translation check between the protein sequence meta data outputted from extract_genomic_information_from_uniprot_id and the translation of the genomic loci exons
    outputted from this function 

    Args:
    genomic_information (pd.DataFrame): dataframe input from the output of extract_genomic_information_from_uniprot_id

    Returns:
    genomic_information_loci_extracted (pd.DataFrame): dataframe output with extracted loci, loci_exons and translation checks
    '''
    
    #print(genomic_information)
    chromosome_id = str(genomic_information['gnCoordinate.genomicLocation.chromosome'].iloc[0])
    records_dict = {seq.id: seq for seq in genome_sequence_records}
    match = records_dict.get(chromosome_id)
    if match:
        is_reverse_strand = str(genomic_information['gnCoordinate.genomicLocation.reverseStrand'].iloc[0]) == 'True'
        uniprot_start = int(genomic_information['gnCoordinate.genomicLocation.start'].iloc[0])
        uniprot_end = int(genomic_information['gnCoordinate.genomicLocation.end'].iloc[0])
        
        # Adjust start and end for reverse strand
        if is_reverse_strand:
            uniprot_start, uniprot_end = uniprot_end, uniprot_start
           
        # Preparing exons DataFrame by splitting on commas
        df_exons_id = genomic_information[['exon_id']].copy()
        df_exons_id['exon_id'] = df_exons_id['exon_id'].str.split(',')
        
        df_exons_start = genomic_information[['exon_genomeLocation.begin.position']].copy()
        df_exons_start['exon_genomeLocation.begin.position'] = df_exons_start['exon_genomeLocation.begin.position'].str.split(',')
        
        df_exons_end = genomic_information[['exon_genomeLocation.end.position']].copy()
        df_exons_end['exon_genomeLocation.end.position'] = df_exons_end['exon_genomeLocation.end.position'].str.split(',')
        
        # Exploding columns
        df_exons_id = df_exons_id.explode('exon_id')
        df_exons_start = df_exons_start.explode('exon_genomeLocation.begin.position')
        df_exons_end = df_exons_end.explode('exon_genomeLocation.end.position')
        
        df_exons = pd.concat([df_exons_id, df_exons_start, df_exons_end], axis=1)
        df_exons.columns = ['exon_id', 'exon_start', 'exon_end']
        
        if is_reverse_strand:
            df_exons['exon_start'], df_exons['exon_end'] = df_exons['exon_end'], df_exons['exon_start']
        
        extracted_gene = match.seq[(uniprot_start - 1) - UTR_length:(uniprot_end - 1) + UTR_length].lower()
        extracted_gene = str(extracted_gene)
        #print(f'Extracted gene (length:{len(extracted_gene)}):')
        #print(extracted_gene)

        # Convert each exon sequence to uppercase in the extracted gene sequence
        for _, row in df_exons.iterrows():
            if row['exon_start'] != 'nan' and row['exon_end']!= 'nan':
                exon_start = int(float(row['exon_start'])) - uniprot_start + UTR_length
                #print("Exon start:")
                #print(exon_start)
                exon_end = int(float(row['exon_end'])) - uniprot_start + UTR_length
                #print("Exon end:")
                #print(exon_end)
            else:
                exon_start = exon_end = 0
                
            extracted_exon = extracted_gene[exon_start:exon_end+1]
            #print('Extracted exon:')
            #print(extracted_exon)
            
            extracted_gene = (
                    extracted_gene[:exon_start]
                    + extracted_exon.upper()
                    + extracted_gene[exon_end+1:]
                )
            
            #print('Extracted gene:')
            #print(extracted_gene)
        
        if is_reverse_strand:
            annotated_gene = reverse_complement(extracted_gene)
        else:
            annotated_gene = extracted_gene
        
        genomic_loci, genomic_loci_exons = clean_loci(annotated_gene)
        genomic_information_loci_extracted = genomic_information.assign(genomic_loci=genomic_loci, genomic_loci_exon=genomic_loci_exons)
        
        #Print reference loci and amino acid sequence 
        #print(f'Genomic loci exons (length = {len(genomic_loci_exons)})')
        #print(genomic_loci_exons)
        #print(f'Reference amino acid sequence (length = {len(genomic_information_loci_extracted['sequence'].iloc[0])}):')
        #print(genomic_information_loci_extracted['sequence'].iloc[0])

        # Translate the extracted loci exons and match them against the uniprot sequence 
        if genomic_loci_exons.startswith("ATG"):
            translation = translate(genomic_loci_exons)
            #print('Translating exons...')
            #print(translation)
            if translation == genomic_information_loci_extracted['sequence'].iloc[0]:
                genomic_information_loci_extracted = genomic_information_loci_extracted.assign(translation_check='Pass')
                print('Translation matches protein sequence')
                return genomic_information_loci_extracted
            else:
                genomic_information_loci_extracted = genomic_information_loci_extracted.assign(translation_check='Fail')   
                print('Translation does not match protein sequence')
                print(genomic_information_loci_extracted)
                return genomic_information_loci_extracted
        else:
            genomic_information_loci_extracted = recursive_translate(genomic_loci_exons, genomic_information_loci_extracted)
            return genomic_information_loci_extracted

# %% [code]
#Load up reference genome, keep this in the same folder as the script
test_multipart_fasta = 'Homo_sapiens.GRCh38.dna.toplevel.fa'
test_sequence_records = extract_genome_multifast_to_list_of_sequence_records(test_multipart_fasta)
print(test_sequence_records)


# %% [code]
#for batch 
pd.set_option('display.max_columns', None)

#Enter_excel_sheet_name_here
test_uniprot_id_dataframe = pd.read_excel('uniprotkb_AND_model_organism_9606_AND_r_2025_05_02.xlsx') #Change this for excel sheet name. 2 Column format ['Uniprot_ID'],[Amino_acid_position]
display(test_uniprot_id_dataframe)

#Identify unique uniprot IDs for uniprot API search and subsequent loci extraction
test_uniprot_id_dataframe_unique = pd.DataFrame()
test_uniprot_id_dataframe_unique['Uniprot_ID'] = test_uniprot_id_dataframe['Uniprot_ID'].unique()
display(test_uniprot_id_dataframe_unique)

#Using iterrows to apply the function extract_genomic_information_from_uniprot_id
test_genomic_information_list = []
extraction_failure_list = []
for index, row in test_uniprot_id_dataframe_unique.iterrows():

    extracted_genomic_information,extraction_failure = extract_genomic_information_from_uniprot_id(row['Uniprot_ID'])
    print(extraction_failure)
    test_genomic_information_list.append(extracted_genomic_information)
    extraction_failure_list.append(extraction_failure)
    

# print('Lists')
# print(test_genomic_information_list)
# print('Lists')
# print(extraction_failure_list)

test_genomic_information_batch= pd.concat(test_genomic_information_list, ignore_index=True)
display(test_genomic_information_batch)
test_genomic_information_batch.to_excel('Batch_to_trim.xlsx', index = False)

'''
Batch_to_trim file will be generated. remove any duplicates that arise. Keeping the first one of each duplicate usually works.
Save the Batch_to_trim files as Batch_trimmed after duplicates are removed. 
Save it in the same folder as where the script is hosted.
Then move to the next script.
'''


# %% [code]
#Run the Batch_trimmed file 
test_genomic_information_batch_trimmed = pd.read_excel('Batch_trimmed.xlsx').reset_index(drop=True)
display(test_genomic_information_batch_trimmed)

#Using iterrows to apply the function extract_genomic_loci_from_genomic_information
test_genomic_loci_extracted_list = []
i=1
for index, row in test_genomic_information_batch_trimmed.iterrows():
    print(f'{i}/{len(test_genomic_information_batch_trimmed)}')
    test_genomic_loci_extracted_list.append(extract_genomic_loci_from_genomic_information(pd.DataFrame([row]), test_sequence_records))
    i+=1
    
test_genomic_loci_extracted_batch = pd.concat(test_genomic_loci_extracted_list, ignore_index=True)
display(test_genomic_loci_extracted_batch)

# test_genomic_loci_extracted_batch_plus_aa= pd.merge(test_uniprot_id_dataframe, test_genomic_loci_extracted_batch, left_on='Uniprot_ID', right_on = 'accession', how='left')
# display(test_genomic_loci_extracted_batch_plus_aa)

#Note test genomic_loci_extracted_batch_plus_aa needs to be kept in memory as opening in excel will cut off the character count for things like the loci.

# %% [code]
#Get statistics for figure
#Total number of inputs
total_human_swiss_prot = 20421
print('Total Swiss_prot:')
print(total_human_swiss_prot)

#Numbers passed through to genomic extraction 
total_passed_through_to_genomic_extraction = 19116
print('Total Passed to genomic extraction:')
print(total_passed_through_to_genomic_extraction)

#Numbers passed translations check
total_passed_by_translation_check = (test_genomic_loci_extracted_batch['translation_check'] == 'Pass').sum()
print('Total Passed by translation check:')
print(total_passed_by_translation_check)


# %% [code]
# For custom loci e.g for anything that doesn't have a good reference genome on uniprot

def custom_loci(input_dataframe): 
    display(input_dataframe)
    print()
    annotated_gene = str(input_dataframe.at[0, 'genomic_loci'])
    genomic_loci, genomic_loci_exons = clean_loci(annotated_gene)
    genomic_information_loci_extracted = input_dataframe.assign(genomic_loci = genomic_loci, genomic_loci_exon = genomic_loci_exons)
        
    #Print reference loci and amino acid sequence 
    print(f'Genomic loci exons (length = {len(genomic_loci)})')
    print(genomic_loci)
    print(f'Genomic loci exons (length = {len(genomic_loci_exons)})')
    print(genomic_loci_exons)
    print(f'Reference amino acid sequence (length = {len(genomic_information_loci_extracted['protein_sequence'].iloc[0])}):')
    print(genomic_information_loci_extracted['protein_sequence'].iloc[0])
    
    # Translate the extracted loci exons and match them against the uniprot sequence 
    if genomic_loci_exons.startswith("ATG"):
        translation = translate(genomic_loci_exons)
        print('Translating exons...')
        print(translation)
        if translation == genomic_information_loci_extracted['protein_sequence'].iloc[0]:
            genomic_information_loci_extracted = genomic_information_loci_extracted.assign(translation_check='Pass')
            print('Translation matches protein sequence:')
            return genomic_information_loci_extracted
        else:
            genomic_information_loci_extracted = genomic_information_loci_extracted.assign(translation_check='Fail')   
            print('Translation does not match protein sequence (running string search as possible fix):')
            return genomic_information_loci_extracted
    else:
        genomic_information_loci_extracted = recursive_translate(genomic_loci_exons, genomic_information_loci_extracted)
        return genomic_information_loci_extracted

# %% [code]


# %% [code]
#Custom loci handling

# test_genomic_information_custom_loci = pd.read_excel('custom_loci_tests.xlsx').reset_index(drop=True)
# display(test_genomic_information_custom_loci)

# print(test_genomic_information_custom_loci['genomic_loci'])

# print(test_genomic_information_custom_loci.at[0, 'genomic_loci'])

# #Using iterrows to apply the function extract_genomic_loci_from_genomic_information
# test_genomic_information_custom_loci_list = []
# for index, row in test_genomic_information_custom_loci.iterrows():
#     display(row)
#     test_genomic_information_custom_loci_list.append(custom_loci(pd.DataFrame([row])))
#     break
                                            
# test_genomic_loci_extracted_batch = pd.concat(test_genomic_information_custom_loci_list, ignore_index=True)
# display(test_genomic_information_custom_loci_list)

# %% [code]
# guide generation for specific amino acid
def find_guides(loci, pam):
    """ 
    Finds guide RNA sequences on the positive and negative strands of the loci by 
    searching for PAM sites and extracting 20 bases upstream from the PAM position. 
    
    Arguments:
        loci - DNA input sequence as a string
        pam - protospacer adjacent motif for Cas protein
        
    Returns:
        crRNA_DNA_start_position - position of crRNA_DNA start site (adjusted for python 0 indexing)
        crRNA_DNA_sequence - sequence 20bp upstream of PAM
        strand - which strand PAM was found on (positive or negative)
    """
    
    # Initialize lists to store gRNA information
    pam_found = []
    crRNA_DNA_sequence_start_position = []
    crRNA_DNA_sequence = []
    strand = []
    
    # Define PAM motifs
    if pam == "NGG":
        print(f'Searching forward strand for {pam} sites...')
        print(loci)
        for n in range(len(loci) - 2):  # Ensure n + 2 in the following line is within bounds
            if (loci[n] in ['A','a','T','t','C','c','G','g'] and loci[n+1] in ['G','g'] and loci[n+2] in ['G','g']):
                if n - 21 >= 0:  # Ensure there are enough bases before the PAM
                    pam_found.append(loci[n:n+3])
                    crRNA_DNA_sequence_start_position.append(n) # crRNA DNA sequence start site
                    crRNA_DNA_sequence.append(loci[n-20:n])  # Extract sequence (adjusted for python 0 indexing and python slicing being non inclusive of the final value)
                    strand.append("forward")
        
        print(f'Searching reverse strand for {pam} sites...')
        loci_reverse_complement = reverse_complement(loci)
        print(loci_reverse_complement)
        for n in range(len(loci_reverse_complement) - 2):  # Ensure n + 2 in the following line is within bounds
            if (loci_reverse_complement[n] in ['A','a','T','t','C','c','G','g'] and loci_reverse_complement[n+1] in ['G','g'] and loci_reverse_complement[n+2] in ['G','g']):
                if n - 21 >= 0:  # Ensure there are enough bases before the PAM
                    pam_found.append(loci_reverse_complement[n:n+3])
                    crRNA_DNA_sequence_start_position.append(len(loci) - (n))  #crRNA DNA sequence start site
                    crRNA_DNA_sequence.append(loci_reverse_complement[n-20:n])  # Extract sequence
                    strand.append("reverse")
                    
    else:
        raise ValueError("PAM not recognised")

    return pam_found, crRNA_DNA_sequence_start_position, crRNA_DNA_sequence, strand

#get codon index function
def get_codon_index(loci, loci_exon):
    """ 
    Maps the positions of the protein coding DNA sequences (exons) along the loci. It gives information 
    regarding the amino acid residue in each position.

    Args:
    loci (str): extracted DNA loci sequence 
    loci_exon (str): extracted DNA loci exon sequence

    Returns:
    protein_dict (dict): a dictionary where each base in the loci exon is mapped to a protein and its location in the loci
    """
    assert len(loci_exon) % 3 == 0, "The length of the coding sequence is not a multiple of 3" #Check that exon is a multiple of 3
    Base_df = pd.DataFrame(list(loci), columns = ["Base"]) #List of all the bases in the genomic loci
    Base_df["Position"] = Base_df.index
    Base_df = Base_df[Base_df['Base'].str.isupper()].reset_index(drop=True) # Only
    pos1 = Base_df[Base_df.index % 3 == 0].reset_index(drop=True) #Generate a list of every third base from the first base
    pos2 = Base_df[Base_df.index % 3 == 1].reset_index(drop=True) #Generate a list of every third base from the second base
    pos3 = Base_df[Base_df.index % 3 == 2].reset_index(drop=True) #Generate a list of every third base from the third base
    aas = list(translate(loci_exon))
    protein_dict = {}
    for aa in range(len(aas)):
        protein_dict[aa] = {"Amino Acid": aas[aa],
                        "base_1": (pos1["Position"][aa],
                                pos1["Base"][aa]),
                        "base_2": (pos2["Position"][aa],
                                pos2["Base"][aa]),
                        "base_3": (pos3["Position"][aa],
                                pos3["Base"][aa])}

    return protein_dict

def get_GC_content(grna_dna_sequence):
    
    """ 
    Return the G/C content of the guide RNA sequence as a percentage.

    args: 
    grna_dna_sequence (str) : dna complement of gRNA variable region

    returns:
    gc_percentage (float): %GC of the dna complement of the gRNA variable region

    """
    gc_count = 0
    base_count = 0
    for char in grna_dna_sequence:
        if char.isalpha(): #Count the length of the guide RNA
            gc_count += 1
        if char in ['C','c','G','g']:
            base_count += 1 #Count the number of Gs or Cs in the guide RNA
    gc_percentage = float(base_count)/float(gc_count) * 100 #Calculate a percentage of Gs and Cs in the length of the guide RNA
    gc_percentage = round(gc_percentage, 2) #Round to two decimal places
    return gc_percentage

def guide_RNA_notes(grna_dna_sequence, gc_percentage):
    """ 
    Return additional information on the genereated guide RNA sequences.
    args:
    grna_dna_sequence (str) : dna complement of gRNA variable region
    gc_percentage (float): %GC of the dna complement of the gRNA variable region
    
    returns:
    notes (list): list of additional information for the guide RNA sequence 

    """

    # Check if guides have been identified
    if grna_dna_sequence == "No 5' guide could be identified" or grna_dna_sequence == "No 3' guide could be identified":
        return ""
    
    # Initialise the notes  
    notes = []
    for n in range(len(grna_dna_sequence)-3):
        if grna_dna_sequence[n:n + 4] == 'TTTT': #Check for four thymines in a row
            notes.append('PolyT present.')
    if grna_dna_sequence[0] != 'G': #Check the guide RNA starts with a 'G' at the most 5' position
        notes.append('No leading G.')
    if gc_percentage >= 75:
        notes.append('G/C content over 75%.') #Check if the G/C content of the guide is more than or equal to 75%
    return notes

def specific_function_guide_RNA_generation(spec_aa_pos, loci, minimum_distance = 5, maximum_distance = 86, pam = 'NGG'):
    """ 
    Identifies and produces metrics for guides upstream and downstream from a specific amino acid residue.
    
    Arguments:
        spec_aa_pos - specific amino acid residue selected by position
        minimum_distance - minimum search distance to find guide RNA (in bases pairs from the codon for the specific amino acid)
        minimum_distance - maximum search distance to find guide RNA (in bases pairs from the codon for the specific amino acid)
        pam - pam sequence e.g. NGG for spCas9
        loci - DNA loci for protein of interest 
            
    Returns:
        guides (pd.Dataframe): pandas dataframe containing information about all guide pairs
    """
  
    loci, loci_exon = clean_loci(loci)
    print(f'loci (length:{len(loci)})')
    print(loci)
    print(f'loci exon (length:{len(loci_exon)})')
    print(loci_exon)
    
    # convert the CDS to a protein dictionary
    protein_dict = get_codon_index(loci, loci_exon)
    
    # display information about the selected amino acid
    spec_aa_pos = int(spec_aa_pos)
    maximum_distance = int(maximum_distance)
    aa_selected = protein_dict[spec_aa_pos - 1] #-1 corrects for python 0 indexing
    # Confirm the selection is the correct amino acid
    print(f'The amino acid you have selected is {aa_selected} at aa position {spec_aa_pos} and loci postion {protein_dict[spec_aa_pos - 1]["base_1"][0]} - {protein_dict[spec_aa_pos - 1]["base_3"][0]}')  
    print(f'The PAM you have selected is {pam}')
    
    # generate lists of all the potential guides within a subsection genomic loci and a list of all guides on the reverse complement.
    loci_region_start = protein_dict[spec_aa_pos - 1]["base_1"][0] - maximum_distance - 25
    loci_region_end = protein_dict[spec_aa_pos - 1]["base_3"][0] + maximum_distance + 25
    loci_search_region = loci[loci_region_start:loci_region_end] # Make a smaller region of the loci for loci off target checking + easier visualisation
    print(f'Searching for gRNAs in loci region {loci_region_start}-{loci_region_end}...')
    
    #loci check region for appending onto output dataframe for easy checks and to be used in the 
    loci_check_region = loci[protein_dict[spec_aa_pos - 1]["base_1"][0] - 1000: protein_dict[spec_aa_pos - 1]["base_3"][0] + 1000]


    # Identify PAMs in the inputted genomic loci
    pam_found , guide_positions, gRNA_list, guide_strands = find_guides(loci_search_region, pam)  
    
    # convert the lists into dataframes
    guide_df = pd.DataFrame()
    
    # Add a column for the pam
    guide_df["PAM"] = pam_found
    
    # Add a column of guide RNA sequences to the dataframe
    guide_df["gRNA variable region sequence"] = gRNA_list  
    
    # Add a column of guide RNA cut site positions to the dataframe
    guide_df["Position on + strand"] = guide_positions + loci_region_start
                        
    # Add a column to strand relative to the inputted genomic loci
    guide_df["Strand"] = guide_strands
    
    # Sort the guide RNAs by their position in the genomic loci
    guide_df = guide_df.sort_values(by=["Position on + strand"]).reset_index(drop=True)
    
    # Stops code if guide dataframe is empty
    assert not guide_df.empty, "No guide RNAs identified in sequence"
    
    print(f'{(len(guide_df))} guides found')
    
   #get the distance of the gRNAs from the start and end base of the amino acid
    start_base = protein_dict[spec_aa_pos - 1]["base_1"][0] # -1 corrects for python 0 index
    print(start_base)
    end_base = protein_dict[spec_aa_pos - 1]["base_3"][0] # -1 corrects for python 0 index
    print(end_base)
    guide_df["Distance from start of Amino Acid (bp)"] = guide_df["Position on + strand"] - start_base
    guide_df["Distance from end of Amino Acid (bp)"] = guide_df["Position on + strand"] - end_base - 1 # -1 ensures that distances are not inclusive of the end base
  
  

    # define a dataframe for if there are no guides within the distance range
    noguides = pd.DataFrame(columns=["loci_region","PAM","gRNA variable region sequence","Position on + strand","Strand","Distance from Amino Acid (bp)"])  
       
    # get the upstream guide RNA dataframe
    upstream_guides = []
    upstream_guides = guide_df.query(f"`Distance from start of Amino Acid (bp)` <= {-minimum_distance}").query(f"`Distance from start of Amino Acid (bp)` >= {-maximum_distance}")
    upstream_guides = upstream_guides.rename(columns={"Distance from start of Amino Acid (bp)": "Distance from Amino Acid (bp)"})
    upstream_guides = upstream_guides.drop("Distance from end of Amino Acid (bp)", axis=1)
    print('upstream_guides')
    print(upstream_guides)
    
    if len(upstream_guides) == 0:
        upstream_guides = noguides
    
    # get the downstream guide RNA dataframe
    downstream_guides = []
    downstream_guides = guide_df.query(f"`Distance from end of Amino Acid (bp)` >= {minimum_distance}").query(f"`Distance from end of Amino Acid (bp)` <= {maximum_distance}")
    downstream_guides = downstream_guides.rename(columns={"Distance from end of Amino Acid (bp)": "Distance from Amino Acid (bp)"})
    downstream_guides = downstream_guides.drop("Distance from start of Amino Acid (bp)", axis=1)
    print('downstream_guides')
    print(downstream_guides)
    
    if len(downstream_guides) == 0:
        downstream_guides = noguides
        guides = pd.concat([upstream_guides,downstream_guides])
        return guides

    # join the dataframes together
    guides = pd.concat([upstream_guides,downstream_guides])
    guides = guides.reset_index(drop = True)
    
    # get the G/C content
    guides["G/C Content (%)"] = guides.apply(
        lambda row: (
            get_GC_content(row["gRNA variable region sequence"])
            if row["gRNA variable region sequence"] != ""
            else ""
        ),
        axis=1
    )
    # get the notes
    guides["Notes"] = guides.apply(
        lambda row: (
            guide_RNA_notes(row["gRNA variable region sequence"], row["G/C Content (%)"])
            if row["gRNA variable region sequence"] != ""
            else ""
        ),
        axis=1
    )  
    
    # Perform off target searches
    guides["off target count (loci)"] = guides.apply(
    lambda row: loci.count(row['gRNA variable region sequence'] + row['PAM']) + 
                 loci.count(reverse_complement(row['gRNA variable region sequence'] + row['PAM'])) - 1,  
    axis=1
)
 
      # Add a column for loci_region
    guides["loci_region"] = str(loci_check_region)

    # Reorganise the guide RNA dataframe
    guides = guides[
        [
            "PAM",
            "Distance from Amino Acid (bp)",
            "gRNA variable region sequence",
            "Position on + strand",
            "Strand",
            "G/C Content (%)",
            "off target count (loci)",
            "Notes",
            "loci_region"
        ]
    ]
    guides = guides.reset_index(drop = True)
    
    return guides

#aho corasick search 
def aho_corasick_search(gRNA_sequences, chromosomes):
    """
    Searches for both forward and reverse complement gRNA sequences in a list of Biopython SeqRecord objects using Aho-Corasick.
    Returns a dictionary with total counts of each unique sequence (forward + reverse complement).

    Args:
    gRNA_sequences(str): List of guide RNA sequences.
    chromosomes (list): List of Biopython SeqRecord objects.

    Returns:
    total_matches: Dictionary with combined total counts of each gRNA sequence across all chromosomes.
    """

    # Map each gRNA sequence and it's reverse complement map to a unique key
    print(gRNA_sequences.str.upper())
    print(len(gRNA_sequences))

    print(gRNA_sequences.str.upper().unique())
    print(len(gRNA_sequences.unique()))
    
    sequence_map = {}
    i = 0
    for seq in gRNA_sequences:
        print(f'{i}/{len(gRNA_sequences)}')
        seq = seq.upper()  # Ensure original sequence is uppercase
        rev_seq = reverse_complement(seq)  # Get reverse complement
        
        print(seq)
        print(rev_seq)
        canonical_seq = seq  # Always use `seq` as the canonical representation
        
            # Add `seq` to the dictionary if it is not already present
        if seq not in sequence_map:
            sequence_map[seq] = canonical_seq
        else:
            print(f"Sequence {seq} already exists in the dictionary!")

        # Add reverse complement `rev_seq` to the dictionary if it is not already present
        if rev_seq not in sequence_map:
            sequence_map[rev_seq] = canonical_seq
        else:
            print(f"Reverse complement {rev_seq} already exists in the dictionary!")
        i = i+1

    # Build Aho-Corasick automaton with all unique sequences
    automaton = Automaton()
    for seq in sequence_map.keys():  # Add both forward and reverse sequences
        automaton.add_word(seq, seq)  # Store pattern as value
    automaton.make_automaton()  # Create failure links

    # Initialize dictionary to store total matches (merged forward & reverse complement counts)
    total_matches = {canonical_seq: -1 for canonical_seq in set(sequence_map.values())} # -1 so that the first match sets off target to 0 

    # Search each chromosome separately and sum the counts
    for chrom in chromosomes:
        chrom_seq = str(chrom.seq)  # Convert Seq object to string
        for _, matched_seq in automaton.iter(chrom_seq):
            canonical_seq = sequence_map[matched_seq]  # Map to canonical sequence
            total_matches[canonical_seq] += 1

    total_matches_df = pd.DataFrame(list(total_matches.items()), columns=["Sequence", "off target counts (genome)"])
    return total_matches_df  # Return the final dictionary with merged counts

# %% [code]
#For batch 
#Using iterrows through the DataFrame generated when you ran the first few cells to apply the specific_function_guide_RNA_generation function.
for index, row in test_genomic_loci_extracted_batch_plus_aa.iterrows():
    guides = specific_function_guide_RNA_generation(row['Amino_acid_position'], row['genomic_loci'])
    
    # Now extract gRNA variable sequences and PAM for the aho corasick genome search
    gRNA_sequences = guides['gRNA variable region sequence'] + guides['PAM']
    
    aho_corasick_results = aho_corasick_search(gRNA_sequences, test_sequence_records)
    
    # Merge back into the guides dataframe on full sequence
    # Create a new column for merging
    guides['merged_key'] = guides['gRNA variable region sequence']+ guides['PAM']
    guides['merged_key'] = guides['merged_key'].str.upper()

    # Do another merge on the reverse complement of the aho corasick results because sometimes if two guides are found on the same space on each strand, 
    # then the merge misses a few cases.
    aho_corasick_results['Sequence_reverse_comp'] = aho_corasick_results['Sequence'].apply(reverse_complement)

    # Perform the merge
    guides_1 = guides.merge(
        aho_corasick_results, 
        left_on='merged_key', 
        right_on='Sequence',
        how='left'  # Adjust 'how' as needed (left, right, inner, outer)
    )
    
    guides_2 = guides.merge(
        aho_corasick_results, 
        left_on='merged_key', 
        right_on='Sequence_reverse_comp',
        how='left'  # Adjust 'how' as needed (left, right, inner, outer)
    )

    # Combine the results by filling NaNs from the second merge
    guides = guides_1.combine_first(guides_2)
    
    # Drop the temporary merge key and sequence key from the merge
    guides.drop(columns=['merged_key'], inplace=True)
    guides.drop(columns=['Sequence'], inplace=True)
    guides.drop(columns=['Sequence_reverse_comp'], inplace=True)
    
    print('Guides dataframe:')
    display(guides)
    
    #Save guides file to .csv
    guides.to_csv(f'batch_guides/{row['Uniprot_ID']}_{row['Amino_acid_position']}.csv', index=False) #Guides will be saved in a folder called batch guides. 

# %% [code]
#Function for guide RNA selection
def select_guide_pair(specific_function_output):
    """
    Allows the user to select two rows from a specific function output
    and returns a new DataFrame with the selected rows.
    
    Arguments:
    specific_function_output: The guide RNA DataFrame to select rows from.
    
    Returns:
    upstream guide: row with upstream guide information 
    downstream_guide: row with downstream guide information 
    """

    # make a variable selection_complete to only save the selected guides when certain criteria are met
    selection_complete = False
    # Display the DataFrame
    while selection_complete == False:
        print("Here is the DataFrame of all guides:")
        print("Upstream guides have a negative distance and downstream guides have a positive distance")
        print("If possible, pick a reverse strand gRNA upstream guide and and forward strand downstream guide")
        print("This means that the guide RNA sequence will be recodonised")
        display(specific_function_output)
        
        # Allow the user to select two rows
        row1 = int(input(f"Enter the index of the upstream guide (distance to target aa < 0) [0-{len(specific_function_output)-1}]: "))
        row2 = int(input(f"Enter the index of the downstream guide (distance to target aa > 0) [0-{len(specific_function_output)-1}]: "))

        # Validate that the inputs are within the correct range
        if row1 not in specific_function_output.index or row2 not in specific_function_output.index:
            raise ValueError("Selected index is out of range.")

        # Fetch the selected rows and save them as a new DataFrame
        upstream_guide = specific_function_output.iloc[[row1]].reset_index(drop=True)
        downstream_guide = specific_function_output.iloc[[row2]].reset_index(drop=True)

        hdr_size = int(abs(upstream_guide['Distance from Amino Acid (bp)'])) + int(abs(downstream_guide['Distance from Amino Acid (bp)'])) + 3 # +3 adds target codon
        print(hdr_size)

        if hdr_size < 93:
            selection_complete = True
            print('Guide selection successful. Saving guides')
    
    selected_guides = pd.concat([upstream_guide,downstream_guide]).reset_index(drop=True) 
    selected_guides['hdr_size'] = hdr_size 
          
    return selected_guides

# %% [code]
# Define the folder containing your guide RNA files
folder_path = "batch_guides/"

# Iterate through each file in the folder
for filename in os.listdir(folder_path):
    # Construct the full file path
    print(f'Processing:{filename}')
    filename = str(filename)
    file_path = os.path.join(folder_path, filename)
    # Check if the file is a valid format (e.g., CSV)
    if os.path.isfile(file_path) and filename.endswith('.csv'):  #
        try:
            # Read the file into a DataFrame
            guide_dataframe = pd.read_csv(file_path)  # Change to pd.read_excel() or other methods as needed
            selected_guide_dataframe = select_guide_pair(guide_dataframe)
            display(selected_guide_dataframe)
            selected_guide_dataframe.to_csv(f'batch_selected_guides/selected_guides_{filename}', index = False)
            
        except Exception as e:
            print(f"Error loading {filename}: {e}")

    clear_output()   


    


# %% [code]
#Now set up the code for HDR template generation
def generate_hdr_template(loci, upstream_guide_position, downstream_guide_position, homology_overlap = 60):
    """
    Finds the sequence between two guide RNA sequences in a genomic loci and adds an homology overlap.
    
    Arguments:
    loci (str): Genomic loci that the guides were generated from.
    upstream_guide_position: position of the start of the upstream guide PAM
    downstream_guide_position: position of the start of the downstream guide PAM
    homology_overlap (int): Number of nucleotides to pad around the sequence found between the guides. Default = 40

    Returns:
    hdr_dataframe(pandas DataFrame): Dataframe containing the following information: hdr template sequence, 
    homology arm sequences and their position in the loci

    """
     
    # Get the start and end indices in the loci of the region between guides
    start_hdr_template = int(upstream_guide_position)
    end_hdr_template = int(downstream_guide_position)

        
    # Calculate the indices for the start and end of homology arms
    upstream_homology_arm_start = max(0, start_hdr_template - homology_overlap)
    upstream_homology_arm_end = upstream_guide_position
    downstream_homology_arm_start = downstream_guide_position
    downstream_homology_arm_end = min(len(loci), end_hdr_template + homology_overlap)
        
    # Extract the homology arms
    hdr_template = loci[start_hdr_template:end_hdr_template] #plus one ensures that end hdr template position is included
    upstream_homology_arm = loci[upstream_homology_arm_start:upstream_homology_arm_end]
    downstream_homology_arm = loci[downstream_homology_arm_start:downstream_homology_arm_end]

    # Create a DataFrame using the individual variables
    hdr_dataframe = pd.DataFrame({
    "hdr_template": hdr_template,
    "upstream_homology_arm": upstream_homology_arm,
    "upstream_homology_arm_start": upstream_homology_arm_start,
    "upstream_homology_arm_end": upstream_homology_arm_end,
    "downstream_homology_arm": downstream_homology_arm,
    "downstream_homology_arm_start": downstream_homology_arm_start,
    "downstream_homology_arm_end": downstream_homology_arm_end
    }, index = [0])
    

    return hdr_dataframe

# %% [code]
# Define the folder containing your files
folder_path = "batch_selected_guides/"

# Iterate through each file in the folder
for filename in os.listdir(folder_path):
    # Construct the full file path
    print(f'Processing:{filename}')
    filename = str(filename)
    file_path = os.path.join(folder_path, filename)
    # Check if the file is a valid format (e.g., CSV)
    if os.path.isfile(file_path) and filename.endswith('.csv'):  # Adjust for your file type
        try:
            # Read the file into a DataFrame
            print(file_path)
            selected_guide_dataframe = pd.read_csv(file_path)
            display(selected_guide_dataframe)
            # now extract the correct loci from the loci containing DataFrame generated earlier in the script using the correct part of the file name 
            uniprot_id_from_file_name = str(filename.split('_')[2])
            print(uniprot_id_from_file_name)

            aa_position_from_file_name = str(filename.split('_')[3])
            print(aa_position_from_file_name)
            
            loci = str(test_genomic_loci_extracted_batch_plus_aa[test_genomic_loci_extracted_batch_plus_aa['Uniprot_ID'] == uniprot_id_from_file_name]['genomic_loci'].values[0])
            print(loci)
            
            # extract upstream guide position
            upstream_guide_position = int(selected_guide_dataframe.loc [0, 'Position on + strand'])
            print(upstream_guide_position)

            # extract downstream guide position
            downstream_guide_position = int(selected_guide_dataframe.loc[1, 'Position on + strand'])
            print(downstream_guide_position)

            hdr_template = generate_hdr_template(loci, upstream_guide_position, downstream_guide_position, homology_overlap = 60 )
            display(hdr_template)

            hdr_template.to_csv(f'batch_hdr_templates/hdr_template_{uniprot_id_from_file_name}_{aa_position_from_file_name}', index=False)
            
        except Exception as e:
            print(f"Error loading {filename}: {e}")

# %% [code]
# Generate HDR template library for each hdr template

# def mutate_intronic_guide_sequences(hdr_template):
#     '''
#     Will check the first 5 and last 5 bases in the HDR template and if they are intronic (lower case), it will replace it for a 
#     TTATT sequence. TTATT is selected because of minimal 
#     '''

#     if hdr_template[0:15].islower():
#         hdr_template_intron_mutated = "ttatt" + hdr_template[5:]

#     else:
#         hdr_template_intron_mutated = hdr_template

#     # Check and replace last 5 letters if they are lowercase
#     if hdr_template[-15:].islower():
#         hdr_template_intron_mutated = hdr_template[:-5] + "ttatt"
    
#     else:
#         hdr_template_intron_mutated = hdr_template

#     return hdr_template_intron_mutated


def recodonise_hdr_template(hdr_template, upstream_guide_distance, downstream_guide_distance):
    """
    Recodonise hdr template to codon with nearest amino acid frequency.
    Arguments: 
    hdr_template(str): hdr template sequence to be used used as the template for recodonisation
    upstream_guide_distance (int): distance from start of upstream guide RNA to the start of the codon for the amino acid of interest
    downstream_guide_distance (int): distance from the start of the downstream guide RNA to the end of the codon for the amino acid of interest
    
    Returns: 
    recodonised_hdr_template(str): recodonised template sequence to be used base template for hdr library generation
    spec_aa_codon_start (int): 
    recodonisation_check (str)

    """

    # 3 dictionaries to be used by the function
    # codon_dict pairs each codon to the corresponding amino acid
    codon_dict = {
    'GCT': 'A', 'GCC': 'A', 'GCA': 'A', 'GCG': 'A',
    'CGT': 'R', 'CGC': 'R', 'CGA': 'R', 'CGG': 'R', 'AGA': 'R', 'AGG': 'R',
    'AAT': 'N', 'AAC': 'N',
    'GAT': 'D', 'GAC': 'D',
    'TGT': 'C', 'TGC': 'C',
    'GAA': 'E', 'GAG': 'E',
    'CAA': 'Q', 'CAG': 'Q',
    'GGT': 'G', 'GGC': 'G', 'GGA': 'G', 'GGG': 'G',
    'CAT': 'H', 'CAC': 'H',
    'ATT': 'I', 'ATC': 'I', 'ATA': 'I',
    'TTA': 'L', 'TTG': 'L', 'CTT': 'L', 'CTC': 'L', 'CTA': 'L', 'CTG': 'L',
    'AAA': 'K', 'AAG': 'K',
    'ATG': 'M',  
    'TTT': 'F', 'TTC': 'F',
    'CCT': 'P', 'CCC': 'P', 'CCA': 'P', 'CCG': 'P',
    'TCT': 'S', 'TCC': 'S', 'TCA': 'S', 'TCG': 'S', 'AGT': 'S', 'AGC': 'S',
    'ACT': 'T', 'ACC': 'T', 'ACA': 'T', 'ACG': 'T',
    'TGG': 'W',
    'TAT': 'Y', 'TAC': 'Y',
    'GTT': 'V', 'GTC': 'V', 'GTA': 'V', 'GTG': 'V',
    'TAA': '*', '*': '*', 'TGA': '*'
    }

    #aa_to_codon pairs each amino acid to a codon
    aa_to_codon = {
    'I': ['ATA', 'ATC', 'ATT'], # Isoleucine
    'M': ['ATG'], # Methionine
    'T': ['ACA', 'ACC', 'ACG', 'ACT'], #Threonine
    'N': ['AAC', 'AAT'], # Asparginine 
    'K': ['AAA', 'AAG'], #Lysine
    'S': ['AGC', 'AGT', 'TCA', 'TCC', 'TCG', 'TCT'],  # Serine
    'R': ['AGA', 'AGG', 'CGA', 'CGC', 'CGG', 'CGT'],  # Arginine
    'L': ['CTA', 'CTC', 'CTG', 'CTT', 'TTA', 'TTG'], # Leucine
    'P': ['CCA', 'CCC', 'CCG', 'CCT'], # Proline
    'H': ['CAC', 'CAT'], # Histidine 
    'Q': ['CAA', 'CAG'], # Gluatmine
    'V': ['GTA', 'GTC', 'GTG', 'GTT'], # Valine
    'A': ['GCA', 'GCC', 'GCG', 'GCT'], # Alanine
    'D': ['GAC', 'GAT'], # Aspartic Acid
    'E': ['GAA', 'GAG'], # Glutamic Acid 
    'G': ['GGA', 'GGC', 'GGG', 'GGT'], # Glycine
    'F': ['TTC', 'TTT'], #Phenylalanine
    'Y': ['TAC', 'TAT'], #Tyrosine
    'C': ['TGC', 'TGT'], #Cysteine
    'W': ['TGG'], #Tryptophan
    '*': ['TAA', 'TAG', 'TGA']  # Stop codons
}
    # human frequency dict pairs each codon to a it frequence usage using the codon frequency table found at. 
    human_frequency_dict = {
        # Isoleucine (Ile)
        'ATA': 0.16,
        'ATC': 0.48,
        'ATT': 0.36,
        
        # Methionine (Met)
        'ATG': 1.00,  
        
        # Threonine (Thr)
        'ACA': 0.28,
        'ACC': 0.36,
        'ACG': 0.12,
        'ACT': 0.24,
        
        # Asparagine (Asn)
        'AAC': 0.54,
        'AAT': 0.46,
        
        # Lysine (Lys)
        'AAA': 0.42,
        'AAG': 0.58,
        
        # Serine (Ser)
        'AGC': 0.24,
        'AGT': 0.15,
        
        # Arginine (Arg)
        'AGA': 0.20,
        'AGG': 0.20,
        
        # Leucine (Leu)
        'CTA': 0.07,
        'CTC': 0.20,
        'CTG': 0.41,
        'CTT': 0.13,
        
        # Proline (Pro)
        'CCA': 0.27,
        'CCC': 0.33,
        'CCG': 0.11,
        'CCT': 0.28,
        
        # Histidine (His)
        'CAC': 0.59,
        'CAT': 0.41,
        
        # Glutamine (Gln)
        'CAA': 0.25,
        'CAG': 0.75,
        
        # Arginine (Arg)
        'CGA': 0.11,
        'CGC': 0.19,
        'CGG': 0.21,
        'CGT': 0.08,
        
        # Valine (Val)
        'GTA': 0.11,
        'GTC': 0.24,
        'GTG': 0.47,
        'GTT': 0.18,
        
        # Alanine (Ala)
        'GCA': 0.23,
        'GCC': 0.40,
        'GCG': 0.11,
        'GCT': 0.26,
        
        # Aspartic acid (Asp)
        'GAC': 0.54,
        'GAT': 0.46,
        
        # Glutamic acid (Glu)
        'GAA': 0.42,
        'GAG': 0.58,
        
        # Glycine (Gly)
        'GGA': 0.25,
        'GGC': 0.34,
        'GGG': 0.25,
        'GGT': 0.16,
        
        # Serine (Ser)
        'TCA': 0.15,
        'TCC': 0.22,
        'TCG': 0.06,
        'TCT': 0.18,
        
        # Phenylalanine (Phe)
        'TTC': 0.55,
        'TTT': 0.45,
        
        # Leucine (Leu)
        'TTA': 0.07,
        'TTG': 0.13,
        
        # Tyrosine (Tyr)
        'TAC': 0.57,
        'TAT': 0.43,
        
        # Cysteine (Cys)
        'TGC': 0.55,
        'TGT': 0.45,
        
        # Tryptophan (Trp)
        'TGG': 1.00,
        
        # Stop (*)
        'TAA': 0.61,
        'TAG': 0.09,
        'TGA': 0.30
    }

    
  
    #First check that the hdr_template_exon is in frame and shift start point of recodonisation to be in frame
    #this is done by identifying the target codon and frameshifting to that reference
    hdr_template, hdr_template_exon = clean_loci(hdr_template)
    spec_aa_codon_start = 0 + abs(upstream_guide_distance)
    spec_aa_codon_end = len(hdr_template) - downstream_guide_distance
    
    spec_aa_codon = hdr_template[spec_aa_codon_start:spec_aa_codon_end]
    print(f'hdr template(length:{len(hdr_template)})')
    print(hdr_template)
    
    print(f'hdr template exon (length:{len(hdr_template_exon)})')
    print(hdr_template_exon)
    
    print(f'Target codon is at position: {spec_aa_codon_start} - {spec_aa_codon_end - 1}')
    print(spec_aa_codon)
    
    print('Translation')
    print(f'{translate(spec_aa_codon)}')

    #Correct spec_aa_codon_start and end for possible introns and the start and the end
    exon_start_index = next((i for i, char in enumerate(hdr_template) if char.isupper()), -1)
    spec_aa_codon_start_exon_corrected = spec_aa_codon_start - exon_start_index

    
    if spec_aa_codon_start_exon_corrected % 3 == 0:
        print('Exons are in frame')
        n = 0
    elif spec_aa_codon_start_exon_corrected % 3 == 1:
        print('Exons are out of frame by 1 position. Shifting start of recodonisation by 1')
        n = 1
    elif spec_aa_codon_start_exon_corrected % 3 == 2:
        print('Exons are out of frame by 2 positions. Shifting start of recodonisation by 2')
        n = 2
        
    #changing codons to aa
    aa_sequence = ""
    hdr_template_exon_dict = {}
    exon_start_index = next((i for i, char in enumerate(hdr_template) if char.isupper()), -1)
    for i in range(n, len(hdr_template_exon)-2, 3):
        codon_start = i
        codon_end = i+3
        codon = hdr_template_exon[codon_start:codon_end]
        print(codon)
        aa = codon_dict[codon]
        aa_sequence += aa
        hdr_template_exon_dict[i // 3] = codon
   
    print(f'hdr sequence to be recodonised(length:{len(hdr_template[n::])})')
    print(hdr_template[n::])
    print('aa sequence')
    print(aa_sequence)
    
    recodonised_sequence = ""
    
    
    for m, aa in enumerate(aa_sequence):
        current_codon = hdr_template_exon_dict[m]

        if aa in ['M', "W", "*"]:
            recodonised_sequence += current_codon
        else:
            #print(aa, current_codon)
            aa_dict = {codon : human_frequency_dict[codon] for codon in aa_to_codon[aa]}
            codon_freq_list = sorted(aa_dict.items(), key=lambda x: x[1], reverse=True)
            for i, codon_freq in enumerate(codon_freq_list):
                if (codon_freq[0] == current_codon) and (i == 0):
                    recodonised_sequence += codon_freq_list[1][0]  # lower down a rank
                elif codon_freq[0] == current_codon:
                    recodonised_sequence += codon_freq_list[i-1][0]  # upper a rank
                    
    print(f'recodonised sequence (length:{len(recodonised_sequence)})')
    print(recodonised_sequence)
    
    print('translating recodonised sequence')
    recodonised_sequence_translation = translate(recodonised_sequence)
    print(recodonised_sequence_translation)
    
    if aa_sequence == recodonised_sequence_translation:
        print('Recodonised translations matches original translation')
    else:
        print('Recodonisation failed')
    
    # Now reattach the recodonised hdr region back to the hdr template
    # find region upstream to reattach to recodonised sequence. +n + 1 correct for the earlier frameshift correction
    exon_start_index = next((i for i, char in enumerate(hdr_template) if char.isupper()), -1)
    upstream_hdr_template = hdr_template[0:exon_start_index+n]
    print('Upstream hdr template')
    print(upstream_hdr_template)
    downstream_hdr_template = hdr_template[exon_start_index+n+len(recodonised_sequence)::]
    print('Downstream hdr template')
    print(downstream_hdr_template)
    
    hdr_template_recodonised = upstream_hdr_template + recodonised_sequence + downstream_hdr_template
    print(f'recodonised hdr template(length:{len(hdr_template_recodonised)})')
    print(hdr_template_recodonised)
    
    if len(hdr_template) == len(hdr_template_recodonised):
        print('Length of hdr template = length of recodonised hdr template')
        recodonisation_check = 'Pass'
    else:
        print('hdr template reconstruction failed')
        recodonisation_check = 'Fail'
        
    # return the recodonized sequence
    return hdr_template_recodonised, spec_aa_codon_start

def reconstruct_loci_with_recodonised_hdr_template(loci, hdr_template_recodonised, upstream_homology_arm_end, downstream_homology_arm_start):
    '''
    This function integrates the recodonised hdr template back into the loci. 

    Args:
    loci (str): Genomic loci that the guides were generated from. 
    hdr_template_recodonised (str): hdr template recodonised to be synonymouse to the wild type 
    upstream_homology_arm_end (int): position in the loci of the end of the upstream homology arm
    downstream_homology_arm_start (int): position in the loci of the start of the downstream homology arm

    Returns:
    section_loci_plus_recodonised_hdr_template (str): Section of the loci with the recodonised hdr template integrated. Used for visual inspection and automated primer design
    '''
    upstream_loci = loci[0:int(upstream_homology_arm_end)]
    downstream_loci = loci[int(downstream_homology_arm_start)::]
    
    loci_plus_recodonised_hdr_template = upstream_loci + hdr_template_recodonised + downstream_loci
    print('Full loci with recodonised HDR template integrated:')
    print(loci_plus_recodonised_hdr_template)

    if len(loci) == len(loci_plus_recodonised_hdr_template):
        print(f'Reconstruction successful -> loci ({len(loci)}) equals loci with hdr template integrated ({len(loci_plus_recodonised_hdr_template)})')
        
        section_loci_plus_recodonised_hdr_template = loci_plus_recodonised_hdr_template[int(upstream_homology_arm_end) - 500:int(downstream_homology_arm_start) + 500]
        print('Section of loci with recodonised HDR template integrated:')
        print(section_loci_plus_recodonised_hdr_template)
        
        return loci_plus_recodonised_hdr_template, section_loci_plus_recodonised_hdr_template
    else:
        print(f'Reconstruction failed -> loci ({len(loci)}) not equal to loci with hdr template integrated ({len(loci_plus_recodonised_hdr_template)})')
        return

def design_integration_specific_primers (uniprot_id_amino_acid, hdr_template_recodonised, loci_plus_recodonised_hdr_template, upstream_homology_arm_end, upstream_guide_distance, downstream_homology_arm_start, downstream_guide_distance):
    '''
    This function will design integration specific primers to target redocodonised HDR templates. These primer pairs will be used to generated amplicons for illumina sequencing

    Args:
    uniprot_id_amino_acid (str): ID for primer pair
    hdr_template_recodonised (str): hdr template recodonised to be synonymous to the wild type
    loci_plus_recodonised_hdr_template (str): Full genomic loci with recodonised hdr template included
    upstream_homology_arm_end (int): numerical position of end of upstream homology arm in the loci 
    upstream_guide_distance (int): numerical distance between end of the upstream homology arm and the start of the codon for the amino acid of interest
    downstream_homology_arm_start (int): numerical position of start of the downstream homology arm in the loci 
    downstream_guide_distance (int): numerical distance between start of the upstream homology arm and the end of the codon for the amino acid of interest
    
    Returns:
    integration_specific_primers (DataFrame): dataframe containing forward and reverse primer sequences and Tm's
    '''
    
    #Make sure integration-specific primer falls within the exonic portion of the recodonised region
    exon_start_index = upstream_homology_arm_end
    upstream_primer_range = upstream_guide_distance
    if loci_plus_recodonised_hdr_template[exon_start_index].islower():
        first_upper = next((i for i, char in enumerate(hdr_template_recodonised) if char.isupper()), -1)
        exon_start_index = first_upper + upstream_homology_arm_end
        upstream_primer_range = upstream_guide_distance + first_upper

    exon_end_index = downstream_homology_arm_start
    downstream_primer_range = downstream_guide_distance
    if loci_plus_recodonised_hdr_template[exon_end_index].islower():
        last_upper = next((i for i, char in enumerate(hdr_template_recodonised[::-1]) if char.isupper()), -1)
        exon_end_index = downstream_homology_arm_start - last_upper
        downstream_primer_range = downstream_guide_distance - last_upper

    print("exon start index: " + str(exon_start_index))
    print("exon end index: " + str(exon_end_index))

    #Add logic to make sure that all recodonised region specific primers at at most 50 bp from the codon of interest
    upstream_recodonised_primer_offset = 0
    if abs(upstream_primer_range) > 50:
        upstream_recodonised_primer_offset = abs(upstream_primer_range) - 50

    downstream_recodonised_primer_offset = 0
    if downstream_primer_range > 50:
        downstream_recodonised_primer_offset = downstream_primer_range - 50

    print(loci_plus_recodonised_hdr_template[exon_start_index:exon_end_index])
    
    try: 
        if(len(loci_plus_recodonised_hdr_template) - exon_end_index < 1000):
            downstream_primer_search_length = len(loci_plus_recodonised_hdr_template) - exon_end_index
        else:
            downstream_primer_search_length = 1000

        print("fwd primer start: " + str(exon_start_index + upstream_recodonised_primer_offset))
        print("fwd primer search distance: " + str(abs(upstream_primer_range) - 5 - upstream_recodonised_primer_offset))
        print("rev primer start: " + str(exon_end_index))
        print("rev primer search distance: " + str(downstream_primer_search_length))

        primers = primer3.bindings.design_primers(
        seq_args= {
            'SEQUENCE_ID': str(uniprot_id_amino_acid),
            'SEQUENCE_TEMPLATE': str(loci_plus_recodonised_hdr_template),
            'SEQUENCE_PRIMER_PAIR_OK_REGION_LIST': [
                exon_start_index + upstream_recodonised_primer_offset,
                abs(upstream_primer_range) - 5 - upstream_recodonised_primer_offset,
                exon_end_index,
                downstream_primer_search_length
        ]
        },
        global_args={
            'PRIMER_OPT_SIZE': 20,
            'PRIMER_MIN_SIZE': 15,
            'PRIMER_MAX_SIZE': 25,
            'PRIMER_OPT_TM': 60.0,
            'PRIMER_MIN_TM': 58.0,
            'PRIMER_MAX_TM': 62.0,
            'PRIMER_MIN_GC': 20.0,
            'PRIMER_MAX_GC': 80.0,
            'PRIMER_SALT_MONOVALENT': 50.0,
            'PRIMER_DNA_CONC': 500.0,
            'PRIMER_PRODUCT_SIZE_RANGE': [[450,550]]
        })

        print(primers)
        # If no primers are returned, raise an exception
        if primers['PRIMER_LEFT_NUM_RETURNED'] == 0:
            raise ValueError("No primers found")
        
        left_primer_sequence = primers['PRIMER_LEFT'][0]['SEQUENCE']
        print(left_primer_sequence)
        left_primer_tm = primers['PRIMER_LEFT'][0]['TM']
        print(left_primer_tm)
        left_primer_start = primers['PRIMER_LEFT'][0]['COORDS'][0]
        left_primer_len = primers['PRIMER_LEFT'][0]['COORDS'][1]
        
        right_primer_sequence = primers['PRIMER_RIGHT'][0]['SEQUENCE']
        print(right_primer_sequence)
        right_primer_tm = primers['PRIMER_RIGHT'][0]['TM']
        print(right_primer_tm)
        right_primer_start = primers['PRIMER_RIGHT'][0]['COORDS'][0]
        right_primer_len = primers['PRIMER_RIGHT'][0]['COORDS'][1]

        primer_dataframe = pd.DataFrame({
        "Left primer": left_primer_sequence,
        "Left primer Tm": left_primer_tm,
        "Left primer start": left_primer_start,
        "Left primer length": left_primer_len,
        "Right primer": right_primer_sequence,
        "Right primer Tm": right_primer_tm,
        "Right primer start": right_primer_start,
        "Right primer length": right_primer_len,
        }, index = [0])
    
        return primer_dataframe
    
    except ValueError as e:
        print(e)
        # Handle the case where no primers are found and continue by making overlap primer on the right side
        if(exon_start_index < 1000):
            upstream_search_position = 0
            upstream_primer_search_length = exon_start_index
        else:
            upstream_search_position = exon_start_index - 1000
            upstream_primer_search_length = 1000

        print("fwd primer start: " + str(upstream_search_position))
        print("fwd primer search distance: " + str(upstream_primer_search_length))
        print("rev primer start: " + str(exon_end_index - downstream_primer_range + 5))
        print("rev primer search distance: " + str(downstream_primer_range - 5 - downstream_recodonised_primer_offset))

        primers = primer3.bindings.design_primers(
        seq_args= {
            'SEQUENCE_ID': str(uniprot_id_amino_acid),
            'SEQUENCE_TEMPLATE': str(loci_plus_recodonised_hdr_template),
            'SEQUENCE_PRIMER_PAIR_OK_REGION_LIST': [
                upstream_search_position,
                upstream_primer_search_length,
                exon_end_index - downstream_primer_range + 5,
                downstream_primer_range - 5 - downstream_recodonised_primer_offset
        ]
        },
        global_args={
            'PRIMER_OPT_SIZE': 20,
            'PRIMER_MIN_SIZE': 15,
            'PRIMER_MAX_SIZE': 25,
            'PRIMER_OPT_TM': 60.0,
            'PRIMER_MIN_TM': 58.0,
            'PRIMER_MAX_TM': 62.0,
            'PRIMER_MIN_GC': 20.0,
            'PRIMER_MAX_GC': 80.0,
            'PRIMER_SALT_MONOVALENT': 50.0,
            'PRIMER_DNA_CONC': 500.0,
            'PRIMER_PRODUCT_SIZE_RANGE': [[450,550]]
        })

        print(primers)
        # If no primers are returned, raise an exception
        if primers['PRIMER_LEFT_NUM_RETURNED'] == 0:
            raise ValueError("No integration-specific primer found after target codon")

        left_primer_sequence = primers['PRIMER_LEFT'][0]['SEQUENCE']
        print(left_primer_sequence)
        left_primer_tm = primers['PRIMER_LEFT'][0]['TM']
        print(left_primer_tm)
        left_primer_start = primers['PRIMER_LEFT'][0]['COORDS'][0]
        left_primer_len = primers['PRIMER_LEFT'][0]['COORDS'][1]
        
        right_primer_sequence = primers['PRIMER_RIGHT'][0]['SEQUENCE']
        print(right_primer_sequence)
        right_primer_tm = primers['PRIMER_RIGHT'][0]['TM']
        print(right_primer_tm)
        right_primer_start = primers['PRIMER_RIGHT'][0]['COORDS'][0]
        right_primer_len = primers['PRIMER_RIGHT'][0]['COORDS'][1]

        primer_dataframe = pd.DataFrame({
        "Left primer": left_primer_sequence,
        "Left primer Tm": left_primer_tm,
        "Left primer start": left_primer_start,
        "Left primer length": left_primer_len,
        "Right primer": right_primer_sequence,
        "Right primer Tm": right_primer_tm,
        "Right primer start": right_primer_start,
        "Right primer length": right_primer_len,
        }, index = [0])
    
        return primer_dataframe

def generate_hdr_library(recodonised_hdr_template, spec_aa_codon_start, variant_specific = None, codon_list = [('I', 'ATC'), ('M', 'ATG'), ('T', 'ACC'), ('N', 'AAC'), ('K', 'AAG'), ('S', 'AGC'), ('R', 'CGG'), ('L', 'CTG'), ('P', 'CCC'), ('H', 'CAC'), ('Q', 'CAG'), ('V', 'GTG'), ('A', 'GCC'), ('D', 'GAC'), ('E', 'GAG'), ('G', 'GGC'), ('F', 'TTC'), ('Y', 'TAC'), ('C', 'TGC'), ('W', 'TGG'), ('*', 'TAA')]):
    '''
    Replaces a codon at a given index
    Arguments:
    hdr_template (str) - homologous directed repair template 
    spec_aa_codon_start (int) - start index of codon to be replaced 
    list_of_codons for library (str) - list of codons for library. Default is max frequencies for all 20 amino acids + 1 for stop codon
    variant_specific (str) = nucleotides for variant specific substitution 
    Returns:
    mutated_sequence (str) - sequence with new codon mutated in
    '''
    print(f'Recodonised template (length:{len(recodonised_hdr_template)}')
    print(recodonised_hdr_template)
    
    #mutate sequence on index position
    mutated_sequences = []
    current_codon = recodonised_hdr_template[spec_aa_codon_start:spec_aa_codon_start+3]
    print(f'Codon to be mutated {current_codon} at position {spec_aa_codon_start}-{spec_aa_codon_start+2}')
    current_amino_acid = translate(current_codon)
    print(current_amino_acid)
    
    for aa_codon in codon_list:
        print(aa_codon)
        new_codon = str(aa_codon[1])
        print(new_codon)
        mutated_sequence = recodonised_hdr_template[:spec_aa_codon_start] + new_codon + recodonised_hdr_template[spec_aa_codon_start+3:]
        print(f'mutated_sequence (length:{len(mutated_sequence)})')
        print(mutated_sequence)
        if len(mutated_sequence) == len(recodonised_hdr_template):
            mutated_sequences.append(mutated_sequence)
    
    # Add variant specific codon if it's suppled and not already in the codon list  
    if variant_specific is not None and variant_specific not in codon_list:
        mutated_sequence = recodonised_hdr_template[:spec_aa_codon_start] + variant_specific + recodonised_hdr_template[spec_aa_codon_start+3:]
        mutated_sequences.append(mutated_sequence)

    print(f'hdr template library (size:{len(mutated_sequences)})')
    print(mutated_sequences)
    
    if spec_aa_codon_start % 3 == 0:
        print('Exons are in frame')
        n = 0
    elif spec_aa_codon_start % 3 == 1:
        print('Exons are out of frame by 1 position. Shifting start of recodonisation by 1')
        n = 1
    elif spec_aa_codon_start % 3 == 2:
        print('Exons are out of frame by 2 positions. Shifting start of recodonisation by 2')
        n = 2
    print('hdr template library (amino_acids)')
    
    aa_mutated_sequences = [] 
    for i in mutated_sequences:
        aa_mutated_sequence = translate(i[n::])
        aa_mutated_sequences.append(aa_mutated_sequence)

    print('Translation of mutated sequences:')      
    print(aa_mutated_sequences)
    return mutated_sequences

def generate_genbank_file(uniprot_id_amino_acid, hdr_template_recodonised, loci_plus_recodonised_hdr_template, upstream_homology_arm_end, downstream_homology_arm_start, spec_aa_codon_start, primer_dataframe):
    '''
    Generates a GenBank file for a given locus with recodonised hdr template and integration specific primers
    Arguments:
    uniprot_id_amino_acid (str): ID for primer pair
    hdr_template_recodonised (str): hdr template recodonised to be synonymous to the wild type
    loci_plus_recodonised_hdr_template (str): Full genomic loci with recodonised hdr template included
    primer_dataframe (DataFrame): dataframe containing forward and reverse primer sequences and Tm's

    Returns:
    gb_record (SeqRecord): GenBank object containing recodonised hdr template and primer features
    '''

    exon_start_index = upstream_homology_arm_end
    if loci_plus_recodonised_hdr_template[exon_start_index].islower():
        first_upper = next((i for i, char in enumerate(hdr_template_recodonised) if char.isupper()), -1)
        exon_start_index = first_upper + upstream_homology_arm_end

    exon_end_index = downstream_homology_arm_start
    if loci_plus_recodonised_hdr_template[exon_end_index].islower():
        last_upper = next((i for i, char in enumerate(hdr_template_recodonised[::-1]) if char.isupper()), -1)
        exon_end_index = downstream_homology_arm_start - last_upper

    seq_object = Seq(loci_plus_recodonised_hdr_template)

    gb_record = SeqRecord(seq_object, id=uniprot_id_amino_acid)
    gb_record.annotations["molecule_type"] = "DNA"

    # Add features
    gb_record.features.append(SeqFeature(
        FeatureLocation(upstream_homology_arm_end, downstream_homology_arm_start), type='region_bw_guides'))
    gb_record.features.append(SeqFeature(
        FeatureLocation(exon_start_index, exon_end_index), type='recodonised'))
    gb_record.features.append(SeqFeature(
        FeatureLocation(int(primer_dataframe['Left primer start'][0]), 
                        int(primer_dataframe['Left primer start'][0] + primer_dataframe['Left primer length'][0]),
                        strand=1), type='fwd'))
    gb_record.features.append(SeqFeature(
        FeatureLocation(int(primer_dataframe['Right primer start'][0] - primer_dataframe['Right primer length'][0]+1), 
                        int(primer_dataframe['Right primer start'][0]+1), 
                        strand=-1), type='rev'))
    gb_record.features.append(SeqFeature(
        FeatureLocation(spec_aa_codon_start + upstream_homology_arm_end,
                        spec_aa_codon_start+3 + upstream_homology_arm_end), type='target_codon'))

    return gb_record


# %% [code]
import random
random.seed(1)

# primer list 
primer_sequences = [
'AAACCGGAGCCATACAGTAC',
'AAAGCACTCTTAGGCCTCTG',
'AAAGGGGCCGTCAATATCAG',
'AAATAAGACGACGACCCTCG',
'AACGATGATGCTCACTCTCG',
'AAGAATTACTGACCCCTCGG',
'AAGACGATCCGAGCCATTAC',
'AAGGAACTATGGCATCGAGC',
'AAGGATATGTAGACACCGCC',
'AAGGCCCAGAAGGATACAAC',
'AAGGCGCTCGGATAATACTC',
'AAGGTATGTATAGCGACCGC',
'AATAGGAACCTCTTACGCGG',
'AATATCACGCAAAAGCACCG',
'AATCAGTTTCTTTGGCAGCC',
'AATGCAAAGCTATTAGCGCG',
'AATGCGTCATTTTACACGGC',
'AATGTCCTTAGGCAGTCGTC',
'ACAACGAGCAGACCGAATAG',
'ACAAGGAGTCGGCATATCAC',
'ACAGAACGAACAGGCACTAC',
'ACAGGAAGCAAGGTATACGC',
'ACAGGGTATATTGAGTGCCC',
'ACATAAGCGATCCCAAGGTC',
'ACATCGCATACCAGAACAGG',
'ACATTAAATTTCGCCGTGGC',
'ACCACAGGTCAAGATTCACG',
'ACCCGTATCGCATAAGGATG',
'ACGAGATGATGCACCGATAG',
'ACGATGGGGACATAGAACAC',
'ACGGAGCCCTTATTGTAACC',
'ACGTATGGGGAACACTACAC',
'ACGTGAAACTGTATCGAGCC',
'ACGTTCAGTTTTCCAATGGC',
'ACTAGATTAGCAAGGCACCC',
'ACTGGACCCAATAAAAGGCC',
'ACTTCGATTGGCAAGGACTG',
'AGAACATAGCATTCACGGGG',
'AGACAACAATCTGAGGCTGG',
'AGACAAGCCTTAACCGTAGG',
'AGACACAAGGCTGATTCCAG',
'AGACATGGGATTGACCACAC',
'AGAGAGGCATGATTGACCTC',
'AGAGTTGCACCTAGAATCCG',
'AGATAGATGCTCCGTCAAGC',
'AGATAGTCACGCACAAGACC',
'AGATTAGCCGACTTTCCTGG',
'AGATTAGCTGCCGATACTGG',
'AGATTGTTACTCCGACGGAC',
'AGATTTCCGACGAGATTCCC',
'AGCATCCGTCTAAATCTCGG',
'AGCTATAAGAATTGCCGGGC',
'AGCTATGATCCCGGTGTAAC',
'AGCTCAATCTAACAGTGGGG',
'AGGACACCAGACCAATGAAG',
'AGGGCTAATTACCATCAGCG',
'AGGTGATCTGACGAATGTCC',
'AGTAAAGCATAGTGCCCAGC',
'AGTAGTATCCGAATCGCTGC',
'AGTATCTCAGCAAGGGCAAC',
'AGTATTAGGCGTCAAGGTCC',
'AGTATTCTTACAGCCAGCCG',
'AGTATTGCCGGACTAAACCC',
'AGTCCCAAGTTCAGACGTAC',
'AGTCCGACACAATGTGACAC',
'AGTGAACTGACCGAATCCTC',
'AGTGGTCTGTAAACCGTACC',
'AGTGTTTTCCATTTTCCGCG',
'AGTTATAAGGGTCCGATGCC',
'AGTTGCAGTATCTAACCCGC',
'AGTTGTAATATCACCCGCGC',
'ATACGTGGCTAGCATGAGAC',
'ATACTGTAAGAACCACGCGG',
'ATAGATCATGTCGGCAGTCG',
'ATAGATGGTGCCTACATGCG',
'ATCACAACAAAGGACGGGTC',
'ATCAGACAACACAGAGGCTG',
'ATCCAGGAGGTCTAGGAACC',
'ATCCTAGAAAAGGCGAAGGC',
'ATGCCATGACGACAACTAGC',
'ATGCTAGCTGGAACTATCGG',
'ATTAGGATTGCGAGCGACAC',
'ATTAGTACACTCCGTGAGCG',
'ATTCAAGGGTTGGACGACTC',
'ATTCTCACGACGCAAGATGG',
'ATTGACGGGAACTACACTCG',
'CACTCGATAGGTACAACCGG',
'CAGACCTACGGATCTTAGCG',
'CCACGAGATAAGAGGATGGC',
'CCAGAGCTTAGGGGACATAC',
'CCCGAGGGGAGAAATATACC',
'CCGAGGGAACCATGATACAG',
'CCGGGAGGAAGATATAGCAC',
'CCGGTTGTACCTATCGAGTG',
'CCGTGCGACAAGATTTCAAG',
'CCTTTAACAGGACATGCAGC',
'CGAACGCAAAAGTCCTCAAG',
'CGATAGAACGACCAGGTAGC',
'CGGATCGAACTTAGGTAGCC',
'CGGGAGGAAGTCTTTAGACC',
'CTAATATCCCTGAGCGACGG',
'CTAGGGAACCAGGCTTAACG',
'CTAGGGGATGGTCCAATACG',
'CTATAGAATCCGGGCTGGTC',
'CTGCTAGGGGCTACTTATCG',
'GAAAAGTCCCAATGAGTGCC',
'GAAGTGGTTTGCCTAAACGC',
'GACCATGCAAGGAGAGGTAC',
'GATACATAGACTTGGCCCCG',
'GCACGCAAAAGGACATAACC',
'GCAGCGTTTTAGCCTACAAG',
'GCATAAAGTTGACAGGCCAG',
'GCTAAATAGAGGGAAGCCCC',
'GGAAAACTAAGACAAGGCGC',
'GGAAACAATAACCATCGGCG',
'GGGCACCGATTAAGAAATGC',
'GGGTTGTCTCCTCTGATAGC',
'GTACTCAGAGATTGCCGGAG',
'GTATAAGATCAGCCGGACCC',
'GTATGTCGGCTCTCGTATCG',
'GTTCAGAGGTACGAACCCTC',
'GTTGCATCTAAGCCAAGTGC',
'TAAAGAGAGGGCGTCCAATC',
'TAACGACGTGCCGAACTTAG',
'TAAGATAGCACCACGGATGG',
'TAAGGATTCATCAGGTGCGC',
'TAAGGGACGATGCTTAACCC',
'TACCACGAAATGCACAGGAG',
'TACTGATAATTCGGACGCCC',
'TACTTGAATACCACGTGGCC',
'TAGCCAGGCAAAAGAGATCC',
'TAGCTCGATAATCAAGGGGC',
'TAGTGACCTAATGCCATGGG',
'TAGTTGAGAACACGAACCCG',
'TATAACAGGCTGCTGAGACC',
'TATACTGAAGAACGGCCCAG',
'TATCAATCCGGAACCAGTGC',
'TATCACGGAAGGACTCAACG',
'TCAAAGGAGCACGAACCTAC',
'TCAAGGTCCGTTATGGAACC',
'TCACATAGAAGGACATGGCG',
'TCACTTGGTATCGAGAACGG',
'TCAGCCTTTCATTGATTGCG',
'TCATCGACAAGATACAGGCG',
'TCCAATTATACGGAGCAGGC',
'TCGAATATGCTGTAACCCCG',
'TCGACCAGGTTATCATGAGC',
'TCGAGACAAGAACGATTCCC',
'TCTAGGACTATCACCGGAGG',
'TCTTCATAAGCCAGAGTGCC',
'TCTTGCGATAGACACAAGCC',
'TGAGCCATAAAAGCAAAGCG',
'TGAGCGCAGAACTATCAGAC',
'TGCATAGTATCCCAACAGGG',
'TGCCAAAGGGTAGAGACATC',
'TGCTGAATGAGAAACCTCGG',
'TGGGGACGACTTATAATGCC',
'TGTGGACCCTATCAAACGAG',
'TTAGCTCAGGTCCAAAGTCC',
'TTAGTAGGCAAGCATACCCG',
'TTCGGGAGCGGATTATACAC',
'TTCTGGGACTGGATAACACG',
'TTGACAGACAATCCGTAGGC'
]
print(len(primer_sequences))

# Generate unique pairs
unique_primer_pair_combinations = list(itertools.combinations(primer_sequences, 2))
random.shuffle(unique_primer_pair_combinations)

print(unique_primer_pair_combinations[0])
print(unique_primer_pair_combinations[1])
print(len(unique_primer_pair_combinations))


# %% [code]
# Define the folder containing your files
guide_folder_path = "batch_selected_guides/"
hdr_template_folder_path = "batch_hdr_templates/"

# Iterate through each file in the folder
unique_primer_pair_number = 0
hdr_libraries = {}

for filename in os.listdir(guide_folder_path):
    # Construct the full file path
    print(f'Processing:{filename}')
    filename = str(filename)
    file_path = os.path.join(guide_folder_path, filename)
    # Check if the file is a valid format (e.g., CSV)
    if os.path.isfile(file_path) and filename.endswith('.csv'):  # Adjust for your file type
        try:
            # Read the file into a DataFrame
            print('Opening file at path:')
            print(file_path)
            selected_guide_dataframe = pd.read_csv(file_path)
            print('File DataFrame:')
            display(selected_guide_dataframe)
            
            # now extract the uniprot_id and amino_acid_position from the file name and search it for in the batch hdr_templates folder 
            extract_after_second_underscore = lambda s: s.split('_', 2)[2].removesuffix('.csv') if s.count('_') >= 2 else ''
            uniprot_id_amino_acid_positon = extract_after_second_underscore(filename)
            print('Extracted uniprot ID and amino acid position')
            print(uniprot_id_amino_acid_positon)
            
            # now go through hdr template folder and find the corresponding hdr template filename
            for hdr_filename in os.listdir(hdr_template_folder_path):
                if fnmatch.fnmatch(hdr_filename, f"*{uniprot_id_amino_acid_positon}*"):
                    hdr_file_path = os.path.join(hdr_template_folder_path, hdr_filename)
                    hdr_template_dataframe = pd.read_csv(hdr_file_path)
                    display(hdr_template_dataframe)
            
            # now use the data from the selected guide data frame and the hdr template dataframe to generate the recodonised hdr template
            # extract upstream guide distance
            upstream_guide_distance = int(selected_guide_dataframe.loc [0, 'Distance from Amino Acid (bp)'])
            print('Upstream guide position:')
            print(upstream_guide_distance)

            # extract downstream guide distance
            downstream_guide_distance = int(selected_guide_dataframe.loc[1, 'Distance from Amino Acid (bp)'])
            print('Downstream guide position:')
            print(downstream_guide_distance)

            # extract hdr template 
            hdr_template = str(hdr_template_dataframe.loc[0,'hdr_template'])
            print('HDR template:')
            print(hdr_template)
            
            # now run the intron mutation and hdr recodonisation of the base hdr template
            hdr_template_recodonised, spec_aa_codon_start = recodonise_hdr_template(hdr_template, upstream_guide_distance, downstream_guide_distance)
            print('HDR template recodonised:')
            print(hdr_template_recodonised)

            # now extract features needed to reconstruct the loci with the recodonised HDR template
            uniprot_id_from_file_name = str(filename.split('_')[2])
            loci = str(test_genomic_loci_extracted_batch_plus_aa[test_genomic_loci_extracted_batch_plus_aa['Uniprot_ID'] == uniprot_id_from_file_name]['genomic_loci'].values[0])
            print('Loci:')
            print(loci) 
            
            upstream_homology_arm_end = int(hdr_template_dataframe.loc[0,'upstream_homology_arm_end'])
            print('Upstream homology arm end position:')
            print(upstream_homology_arm_end)

            downstream_homology_arm_start = int(hdr_template_dataframe.loc[0,'downstream_homology_arm_start'])
            print('downstream homology arm position:')
            print(downstream_homology_arm_start)

            # now reconstruct the loci with the recodonised hdr template
            loci_plus_recodonised_hdr_template, section_loci_plus_recodonised_hdr_template = reconstruct_loci_with_recodonised_hdr_template(loci,hdr_template_recodonised,upstream_homology_arm_end,downstream_homology_arm_start)
            
            print('Reconstructing loci with the recodonised HDR template:')
            print(section_loci_plus_recodonised_hdr_template)

            # save section_loci_plus_recodonised_hdr_template and recodonised hdr template to csv
            recodonised_loci_dataframe = pd.DataFrame({
            "Loci with recodonised HDR template integrated": section_loci_plus_recodonised_hdr_template,
            "Recodonised HDR template": hdr_template_recodonised,
            "Spec_aa_codon_start": spec_aa_codon_start
            }, index = [0])

            recodonised_loci_dataframe.to_csv(f'batch_recodonised_locis/{uniprot_id_amino_acid_positon}_recodonised_loci.csv', index = False)
    
            # now design integration specific primers 
            primers = design_integration_specific_primers (uniprot_id_amino_acid_positon, hdr_template_recodonised, loci_plus_recodonised_hdr_template, upstream_homology_arm_end, upstream_guide_distance, downstream_homology_arm_start, downstream_guide_distance)
            print('Integration specific primers selected')
            display(primers)
            
            #save integration specific primers
            primers.to_csv(f'batch_integration_specific_primers/{uniprot_id_amino_acid_positon}_integration_specific_primers.csv', index = False)

            #generate and save a genbank file
            gb_record = generate_genbank_file(uniprot_id_amino_acid_positon, hdr_template_recodonised, loci_plus_recodonised_hdr_template, upstream_homology_arm_end, downstream_homology_arm_start, spec_aa_codon_start, primers)
            SeqIO.write(gb_record, f'batch_genbank_files/{uniprot_id_amino_acid_positon}.gb', 'genbank')
            print('GenBank files generated')

        except Exception as e:
                print(f"Error loading {filename}: {e}")
                


# %% [code]
# now generate the hdr library
hdr_template_folder_path = "batch_hdr_templates/"
recodonised_loci_folder_path= "batch_recodonised_locis/"

for filename in os.listdir(hdr_template_folder_path):
    try:
        # Construct the full file path
        print(f'Processing:{filename}')
        filename = str(filename)
        file_path = os.path.join(hdr_template_folder_path, filename)
        hdr_template_dataframe = pd.read_csv(hdr_file_path)
        display(hdr_template_dataframe)

        # now extract the uniprot_id and amino_acid_position from the file name and search it for in the batch_recodonised_locis folder 
        extract_after_second_underscore = lambda s: s.split('_', 2)[2].removesuffix('.csv') if s.count('_') >= 2 else ''
        uniprot_id_amino_acid_positon = extract_after_second_underscore(filename)
        print('Extracted uniprot ID and amino acid position:')
        print(uniprot_id_amino_acid_positon)

        for filename in os.listdir(recodonised_loci_folder_path):
            if fnmatch.fnmatch(filename, f"*{uniprot_id_amino_acid_positon}.csv"):
                        recodonised_loci_file_path = os.path.join(recodonised_loci_folder_path, filename)
                        print(recodonised_loci_file_path)
                        recodonised_loci_dataframe = pd.read_csv(recodonised_loci_file_path)
                        display(recodonised_loci_dataframe)

        hdr_template_recodonised = recodonised_loci_dataframe.loc[0,'Recodonised HDR template']
        spec_aa_codon_start = recodonised_loci_dataframe.loc[0,'Spec_aa_codon_start']
        print(hdr_template_recodonised)
        
        hdr_library = generate_hdr_library(hdr_template_recodonised, spec_aa_codon_start)
        print('HDR library:')
        print(hdr_library)
            
        print('First sequence in HDR library')
        print(hdr_library[0])

        hdr_library_lengths = [len(sequence) for sequence in hdr_library]
        print('HDR library lengths:')
        print(hdr_library_lengths)

        #extract the upstream and downstream homology arms and append onto the hdr library
        upstream_homology_arm = str(hdr_template_dataframe.loc[0,'upstream_homology_arm'])
        print('Upstream homology arm:')
        print(upstream_homology_arm)
        upstream_homology_arm_length = len(upstream_homology_arm)
        print('Upstream homology arm length:')
        print(upstream_homology_arm_length)
        downstream_homology_arm = str(hdr_template_dataframe.loc[0,'downstream_homology_arm'])
        print('Downstream homology arm:')
        print(downstream_homology_arm)
        downstream_homology_arm_length = len(downstream_homology_arm)
        print('Downstream homology arm length:')
        print(downstream_homology_arm_length)

        #adding upstream homology arm 
        hdr_library_plus_upstream_homology_arm = [upstream_homology_arm + sequence for sequence in hdr_library]
            
        #adding downstream homology arm 
        hdr_library_plus_homology_arms = [sequence + downstream_homology_arm for sequence in hdr_library_plus_upstream_homology_arm]
        print('HDR library plus homology arms:')
        print(hdr_library_plus_homology_arms) 
        hdr_library_plus_homology_arm_lengths = [len(sequence) for sequence in hdr_library_plus_homology_arms]
        print('HDR library plus homology arm lengths:')
        print(hdr_library_plus_homology_arm_lengths)

        #add a length check to check if everything has appended properly
        hdr_library_length_number = 0
        for sequence in hdr_library_plus_homology_arms:
            if len(sequence) == hdr_library_lengths[hdr_library_length_number] + upstream_homology_arm_length + downstream_homology_arm_length:
                print(f'Homology arm attachment successful for sequence:{sequence}')
            else:
                print(f'Homology arm attachment failed for sequence:{sequence}')
            
            hdr_library_length_number=hdr_library_length_number+1  

        # Now add a unique primer pair to the start and end of the library
        pool_specific_primer_pair = unique_primer_pair_combinations[unique_primer_pair_number]
        print(f'Pool specific primer pair {unique_primer_pair_number} :')
        print(pool_specific_primer_pair)

        # adding upstream pool specific primer sequences
        upstream_pool_specific_primer  = unique_primer_pair_combinations[unique_primer_pair_number][0]
        hdr_library_plus_upstream_primers = [upstream_pool_specific_primer + sequence for sequence in hdr_library_plus_homology_arms]
        upstream_primer_length = len(upstream_pool_specific_primer)
        print('upstream_primer_length:')
        print(upstream_primer_length)

        # adding downstream pool specific primer sequences
        downstream_pool_specific_primer = reverse_complement(unique_primer_pair_combinations[unique_primer_pair_number][1])
        downstream_primer_length = len(downstream_pool_specific_primer)
        print('downstream_primer_length:')
        print(downstream_primer_length)
            
        hdr_library_plus_pool_specific_primers = [sequence + downstream_pool_specific_primer  for sequence in hdr_library_plus_upstream_primers]
        print('HDR library plus pool specific primers:')
        print(hdr_library_plus_pool_specific_primers) 
        hdr_library_plus_pool_specific_primers_lengths = [len(sequence) for sequence in hdr_library_plus_pool_specific_primers]
        print('HDR library plus pool specific primer lengths:')
        print(hdr_library_plus_pool_specific_primers_lengths)

        # move to the next unique primer pair for the next iteration of the loop
        unique_primer_pair_number = unique_primer_pair_number + 1

        # add a length check for correct addition of primers
        hdr_library_plus_homology_arm_length_number = 0
        for sequence in hdr_library_plus_pool_specific_primers:
            if len(sequence) == hdr_library_plus_homology_arm_lengths[hdr_library_plus_homology_arm_length_number] + upstream_primer_length + downstream_primer_length:
                print(f'Pool specific primers attachment successful for sequence:{sequence}')
            else:
                print(f'Pool specific primers attachment failed for sequence:{sequence}')
                
            hdr_library_plus_homology_arm_length_number = hdr_library_plus_homology_arm_length_number + 1  
            
        # Repeat for the addition of the bulk primers
        upstream_bulk_primer = 'AAACACGTGGCAAACATTCC' # primer 1 from https://doi.org/10.1093/synbio/ysx008
        print('Upstream bulk primer:')
        print(upstream_bulk_primer)

        upstream_bulk_primer_length = len(upstream_bulk_primer)
        print('Upstream bulk primer length:')
        print(upstream_bulk_primer_length)

        downstream_bulk_primer = 'AAGGACTGCATACCAGGTTG' # primer 14 from https://doi.org/10.1093/synbio/ysx008
        downstream_bulk_primer = reverse_complement(downstream_bulk_primer)
        print('Downstream bulk primer:')
        print(downstream_bulk_primer)

        downstream_bulk_primer_length = len(downstream_bulk_primer)
        print('Downstream bulk primer length:')
        print(downstream_bulk_primer_length)
            
        # add bulk upstream primer
        hdr_library_plus_bulk_upstream_primers = [upstream_bulk_primer + sequence for sequence in hdr_library_plus_pool_specific_primers]
            
        # add bulk downstream primer
        hdr_library_final = [sequence + downstream_bulk_primer for sequence in hdr_library_plus_bulk_upstream_primers]

        # add a length check for successful bulk primer appending plus a check that the final sequences are less than 300 bases
        hdr_library_plus_pool_specific_primers_lengths_number = 0
        for sequence in hdr_library_final:
            if len(sequence) == hdr_library_plus_pool_specific_primers_lengths[hdr_library_plus_pool_specific_primers_lengths_number] + upstream_bulk_primer_length + downstream_bulk_primer_length and len(sequence) <= 300:
                print(f'Bulk primers attachment successful for sequence:{sequence}')
            else:
                 print(f'Bulk primers attachment failed for sequence:{sequence}')
            
            hdr_library_plus_pool_specific_primers_lengths_number = hdr_library_plus_pool_specific_primers_lengths_number + 1  

        # Now save to a dictionary
        hdr_libraries.setdefault(uniprot_id_amino_acid_positon, []).extend(hdr_library_final)
        print('Final hdr library:')
        print(hdr_libraries)
            
    except Exception as e:
         print(f"Error loading {filename}: {e}")

# Convert dictionary into a list of tuples (key, value)
rows = [(key, value) for key, values in hdr_libraries.items() for value in values]

# Create DataFrame
hdr_libraries = pd.DataFrame(rows, columns=['name', 'sequence'])
display(hdr_libraries)

# %% [code]


# %% [code]
```
