## Supplementary_folder_1 for "CORe Designer: a CRISPR design tool for proteome engineering": Figure_2b.ipynb

```
# %% [code]
import pandas as pd 

summary_stats = pd.read_excel('2b_summary_stats.xlsx')
display(summary_stats)

# %% [code]
summary_stats_calculated = pd.DataFrame()
summary_stats_calculated['Name'] = summary_stats['Name']
summary_stats_calculated['No associated genomic coordinates'] = summary_stats['Pre Uniprot extraction'].astype(int) - summary_stats['Post Uniprot extraction'].astype(int)
summary_stats_calculated['Loci extraction successes'] = summary_stats['Post genomic extraction'].astype(int)
summary_stats_calculated['Loci extraction failures'] = summary_stats ['Post Uniprot extraction'].astype(int) - summary_stats['Post genomic extraction'].astype(int)

display(summary_stats_calculated)

# %% [code]
import plotly.express as px

pastel_colors = ["#fc8d59", "#ffffbf", "#91cf60"]  # Light red, light orange, light green


fig = px.bar(
    summary_stats_calculated,
    x="Name",
    y=[
        "No associated genomic coordinates",
        "Loci extraction failures",
        "Loci extraction successes"
    ],
    color_discrete_sequence=pastel_colors
)

fig.update_layout(
    barmode='stack',
    xaxis_tickangle=-45,
    plot_bgcolor='white',      # Background behind the bars
    paper_bgcolor='white',     # Background around the entire figure
    title = 'Swiss-Prot assessment of CORe Designer loci extraction',
    xaxis_title='Organism',
    yaxis_title='Count'
    ),

fig.update_yaxes(tickformat=".0f")
fig.show()

# Export as PDF
fig.write_image("Figure1b.pdf")  # Saves to the current directory


# %% [code]
import seaborn as sns
import matplotlib.pyplot as plt


# 2. Publication styling
sns.set_theme(style="ticks", context="paper", font_scale=1.2)
fig, ax = plt.subplots(figsize=(8, 6))

# 3. Plotting the stacks
# We stack them by defining the 'bottom' parameter for each subsequent layer
# Colors: 0.1 (darkest), 0.5 (medium), 0.9 (lightest)
p1 = ax.bar(summary_stats_calculated['Name'], summary_stats_calculated
            ['Loci extraction successes'], 
            color='0.5', edgecolor='black', label='Successes')

p2 = ax.bar(summary_stats_calculated['Name'], summary_stats_calculated['No associated genomic coordinates'], 
            bottom=summary_stats_calculated['Loci extraction successes'], 
            color='0.3', edgecolor='black', hatch='///', label='No Coordinates')

p3 = ax.bar(summary_stats_calculated['Name'], summary_stats_calculated['Loci extraction failures'], 
            bottom=summary_stats_calculated['Loci extraction successes'] + summary_stats_calculated ['No associated genomic coordinates'], 
            color='0.9', edgecolor='black', hatch='...', label='Failures')

# 4. Aesthetics and Labels
sns.despine()
plt.ylabel("Count", fontweight='bold')
plt.xlabel("Species", fontweight='bold')
plt.title("Loci extraction and annotation results for Swiss-Prot datasets", fontsize=14, pad=20)

# Move legend outside or to an empty area of the plot
plt.legend(title="Extraction Status", loc='upper right', frameon=True)

# 5. Save as PDF for publication
plt.savefig("Figure_2b.pdf", format="pdf", bbox_inches="tight")

plt.show()
```
