## Supplementary_folder_1 for "CORe Designer: a CRISPR design tool for proteome engineering": Figure_2d.ipynb

```
# %% [code]
#Data analysis for figure 2d
import pandas as pd
import numpy as np 

# %% [code]
df = pd.read_excel('figure2d_stats.xlsx')
display(df)

# %% [code]
# Now calculate the means and standard deviations
column_means = df.iloc[:, 3:6].astype(float).mean()
column_std = df.iloc[:, 3:6].astype(float).std()
print(column_means)
print(column_std)


# %% [code]
import plotly.graph_objects as go
# Compute cumulative sum
cumulative_means = column_means.cumsum()
cumulative_means = np.insert(cumulative_means, 0, 0)  # Insert 0 at position 0


column_std = np.insert(column_std, 0, 0)


# Create Plotly figure
fig = go.Figure()

categories = ['start','guides generated', 'hdr library generated', 'primers generated']


# Plot
fig = go.Figure()
fig.add_trace(go.Scatter(
    x=categories,
    y=cumulative_means,
    mode='lines+markers',
    error_y=dict(
        type='data',     
        array=column_std,  
        visible=True,
        thickness=2,
        width=6,
    ),
    line=dict(color="#91cf60", width=3),
    marker=dict(color="#91cf60", size=8)
))


fig.update_layout(
    xaxis_tickangle=0,
    plot_bgcolor='white',      
    paper_bgcolor='white',      
    title = 'CORe Designer runtimes',
    xaxis_title = 'Steps',
    yaxis_title = 'Cumulative time (s)'
)

fig.update_yaxes(tickformat=".0f")

fig.show()
fig.write_image("Figure1d.pdf")  # Saves to the current directory

# %% [code]
import matplotlib.pyplot as plt
import seaborn as sns
import numpy as np

# 1. Set the aesthetic style for publication
sns.set_theme(style="ticks")
sns.set_context("paper", font_scale=1.2) # 'paper' scale is optimized for journals

# Data setup
categories = ['Start', 'Guides generated', 'HDR library generated', 'Primers generated']
# Assuming these were defined in your environment:
# cumulative_means = [0, ...]
# column_std = [0, ...]

# 2. Create the plot
plt.figure(figsize=(8, 6))

# Plot the line and markers
plt.plot(categories, cumulative_means, marker='o', color='black', 
         linewidth=3, markersize=8, label='Runtime')

# Add error bars (Matplotlib's errorbar provides better control for 'publication' style)
plt.errorbar(categories, cumulative_means, yerr=column_std, 
             fmt='none', ecolor = 'black', elinewidth=2, capsize=6)

# 3. Clean up the axes and labels
plt.title('CORe Designer Runtimes', fontsize=14, fontweight='bold', pad=15)
plt.xlabel('Steps', fontsize=12)
plt.ylabel('Cumulative time (s)', fontsize=12)

# Remove the top and right spines (standard publication look)
sns.despine()

# Adjust layout and save
plt.tight_layout()
plt.savefig("Figure_2d.pdf", dpi=300, bbox_inches='tight')
plt.show()

# %% [code]
```
