## Supplementary_folder_1 for "CORe Designer: a CRISPR design tool for proteome engineering": figure_2e.ipynb

```
# %% [code]
# Notebook for figure 1e
import pandas as pd
import plotly.express as px

# %% [code]
#load up excel 
data = pd.read_excel('figure2e_stats.xlsx')
display(data)

# %% [code]
# Create box plot
fig = px.box(data, x='guide_search_length', y='guides_generation')

fig.update_layout(
   xaxis_tickangle=0,
    plot_bgcolor='white',      
    paper_bgcolor='white',      
    title = 'CORe designer runtimes as guide search distance varies',
    xaxis_title = 'Guide search distance',
    yaxis_title = 'Cumulative time (s)'
    )

fig.update_traces(marker_color="#91cf60",    # outlier points
                  line_color="#91cf60",      # outline
                  fillcolor="#ffffff")       # fill
# Show the plot
fig.show()
fig.write_image("Figure1e.pdf")  # Saves to the current directory

# %% [code]
import seaborn as sns
import matplotlib.pyplot as plt

# Set the publication style
sns.set_theme(style="ticks")
plt.figure(figsize=(8, 6))

# Create the box plot
# We use 'color' to set the line colors and 'boxprops' to handle the fill
ax = sns.boxplot(
    data=data, 
    x='guide_search_length', 
    y='guides_generation',
    color="black",           # Sets the default color for lines/whiskers
    linewidth=2,
    fliersize=4,
    boxprops=dict(facecolor="white", edgecolor="black") # Explicitly set fill and edge
)

# Customizing the median line specifically (it often defaults to a different color)
plt.setp(ax.lines, color="black")
# Labels and Title
plt.title('Guide generation runtimes as guide search distance varies', fontsize=14, pad=15)
plt.xlabel('Guide search distance (bases)', fontsize=12)
plt.ylabel('Time for guide generation step (s)', fontsize=12)

# Publication styling
sns.despine()
plt.tight_layout()

# Save high-quality output
plt.savefig("Figure_2e.pdf", bbox_inches='tight')
plt.show()


# %% [code]
```
