## Supplementary_folder_1 for "CORe Designer: a CRISPR design tool for proteome engineering": Figure_3c_data_analysis.ipynb

```
# %% [code]
import pandas as pd
import plotly.graph_objects as go
manual = pd.read_excel('manual_all_reps.xlsx')
display(manual) 

automated = pd.read_excel('auto_all_reps.xlsx')
display(automated)

#drop the rows for All reads and All codons as not needed for analysis 
manual = manual.drop([0,1])
automated = automated.drop([0,1])


# %% [code]
#Calculate means and std of manual and automated
manual_means = manual[['Rep_1_fold_change_wt_normalised', 'Rep_2_fold_change_wt_normalised','Rep_3_fold_change_wt_normalised']].mean(axis=1)
manual_std = manual[['Rep_1_fold_change_wt_normalised', 'Rep_2_fold_change_wt_normalised','Rep_3_fold_change_wt_normalised']].std(axis=1)

automated_means = automated[['Rep_1_fold_change_wt_normalised', 'Rep_2_fold_change_wt_normalised','Rep_3_fold_change_wt_normalised']].mean(axis=1)
automated_std = automated[['Rep_1_fold_change_wt_normalised', 'Rep_2_fold_change_wt_normalised','Rep_3_fold_change_wt_normalised']].std(axis=1)


# %% [code]
import pandas as pd
import seaborn as sns
import matplotlib.pyplot as plt

# 1. Combine data into a single Long-Form DataFrame
data = {
    'Amino Acid': list(manual['label']) + list(automated['label']),
    'Functional score': list(manual_means) + list(automated_means),
    'Method': ['Manual'] * len(manual_means) + ['CORe Designer'] * len(automated_means),
    'Std': list(manual_std) + list(automated_std)
}
df = pd.DataFrame(data)

# 2. Setup Style
sns.set_theme(style="ticks", context="paper", font_scale=1.2)
plt.figure(figsize=(8, 6))

# 3. Plot using Seaborn's native errorbar handling
# We use 'Std' as the source for error bars directly in the function
palette = {"Manual": "white", "CORe Designer": "gray"}
# Ensure we sort the dataframe to match the order Seaborn plotted (Hue-major)
df_sorted = df.sort_values(['Method', 'Amino Acid'], ascending=[False, True])

ax = sns.barplot(
    data= df_sorted, 
    x='Amino Acid', 
    y='Functional score', 
    hue='Method', 
    palette=palette,
    edgecolor='black',
    linewidth=1.5,
    capsize=.1,      # Adds the caps to the error bars
    errorbar=None    # We will overlay specific error bars to avoid size mismatches
)

# 4. The Fail-Safe Error Bar Overlay
# Instead of guessing the patch order, we find the center of each bar 
# and map it to the data directly.
x_coords = [p.get_x() + p.get_width()/2 for p in ax.patches if p.get_height() > 0]

plt.errorbar(
    x=x_coords, 
    y=df_sorted['Functional score'], 
    yerr=df_sorted['Std'], 
    fmt='none', 
    c='black', 
    capsize=4, 
    elinewidth=1.5
)

# Formatting & Legend
plt.title('Comparison of functional scores between manual and CORe Designer methods', fontsize=14)
plt.ylabel('Functional score (fold change normalised to Cys)', fontsize=12)
sns.despine()

# Fix Legend to show hatches
legend = ax.legend(title='Method')


plt.tight_layout()
plt.savefig("figure3c.pdf")
plt.show()

# %% [code]
```
