## Supplementary_folder_1 for "CORe Designer: a CRISPR design tool for proteome engineering": Figure_3d_data_analysis.ipynb

```
# %% [code]
import pandas as pd
import plotly.graph_objects as go
manual = pd.read_excel('manual_all_reps.xlsx')
display(manual) 

automated = pd.read_excel('auto_all_reps.xlsx')
display(automated)

#drop the rows for All reads and All codons as not needed for analysis 
manual = manual.drop([0,1])
automated = automated.drop([0,1])


# %% [code]
#Preprocess both dataframes for plotting
manual_preprocessed = manual.melt(id_vars="label", value_vars=['Rep_1_fold_change_wt_normalised','Rep_2_fold_change_wt_normalised','Rep_3_fold_change_wt_normalised'],
                  var_name="Replicate", value_name="Value")

automated_preprocessed = automated.melt(id_vars="label", value_vars=['Rep_1_fold_change_wt_normalised','Rep_2_fold_change_wt_normalised','Rep_3_fold_change_wt_normalised'],
                  var_name="Replicate", value_name="Value")


# %% [code]
# Create Plotly figure
fig = go.Figure(data=[
    go.Scatter(x=manual_preprocessed['Value'], y=automated_preprocessed['Value'], mode='markers',marker=dict(color="#91cf60", size=8)) 
])


#Set the bar mode to group (side-by-side)
fig.update_layout(
    
    xaxis_title='Functional score for manual',
    yaxis_title='Functional scores for CORe Designer',
    plot_bgcolor='white',      
    paper_bgcolor='white',  
    title='Correlation of functional scores between manual and CORe Designer methods'
)

fig.show()
fig.write_image("figure2e.pdf")  # PDF format

# %% [code]
# Assuming manual_preprocessed and automated_preprocessed have the same length
df = pd.DataFrame({
    "Manual": manual_preprocessed['Value'],
    "Automated": automated_preprocessed['Value'],
    "Label": manual_preprocessed['label']
})

# Create figure
fig = go.Figure()

# Get unique labels
labels = df["Label"].unique()

# Optional: assign colors (Plotly default palette)
import plotly.express as px
colors = px.colors.qualitative.Plotly

# Add one trace per label
for i, label in enumerate(labels):
    df_label = df[df["Label"] == label]
    fig.add_trace(go.Scatter(
        x=df_label["Manual"],
        y=df_label["Automated"],
        mode='markers',
        name=label,
        marker=dict(color=colors[i % len(colors)], size=10)
    ))

# Layout
fig.update_layout(
    title='Correlation of functional scores between manual and CORe Designer methods',
    xaxis_title='Functional score for manual',
    yaxis_title='Functional scores for automated',
    template='plotly_white'
)

fig.show()

# %% [code]
import matplotlib.pyplot as plt
import seaborn as sns

# 1. Set the aesthetic style
sns.set_theme(style="white", palette="deep")
plt.rcParams['font.family'] = 'sans-serif' # Standard for journals

# 2. Initialize the figure
plt.figure(figsize=(8, 6))

# 3. Create the scatter plot
plot = sns.scatterplot(
    x=manual_preprocessed['Value'], 
    y=automated_preprocessed['Value'],
    color= "black",
    s=60,             # Marker size
    edgecolor='w',    # White edge for better separation
    alpha=0.8
)

# 4. Refine Axes and Labels
plt.title('Correlation of Functional Scores: Manual vs. CORe Designer', fontsize=14)
plt.xlabel('Functional score for manual', fontsize=12)
plt.ylabel('Functional scores for CORe Designer', fontsize=12)

# Remove the top and right spines (standard publication look)
sns.despine()

# 5. Save in high-resolution PDF
plt.tight_layout()
plt.savefig("Figure_3d.pdf", format='pdf', dpi=300)
plt.show()

# %% [code]
from scipy.stats import pearsonr
import pandas as pd

# Define replicate columns
replicates = ['Rep_1_fold_change_wt_normalised', 'Rep_2_fold_change_wt_normalised', 'Rep_3_fold_change_wt_normalised']

# Sort by label to align values
manual_sorted = manual.sort_values('label').reset_index(drop=True)
display()
automated_sorted = automated.sort_values('label').reset_index(drop=True)

# Compute correlations for every combination
results = []

for m_col in replicates:
    for a_col in replicates:
        r, p = pearsonr(manual_sorted[m_col], automated_sorted[a_col])
        results.append({
            'Manual': m_col,
            'Automated': a_col,
            'Pearson_r': r,
            'p_value': p
        })

# Convert results to a DataFrame for easy viewing
correlation_df = pd.DataFrame(results)
display(correlation_df)

mean_r = correlation_df['Pearson_r'].mean()
print(f"Mean Pearson r: {mean_r:.3f}")

# %% [code]
```
