## Supplementary figures and images for "CORe Designer: a CRISPR design tool for proteome engineering"

### figure3c.pdf

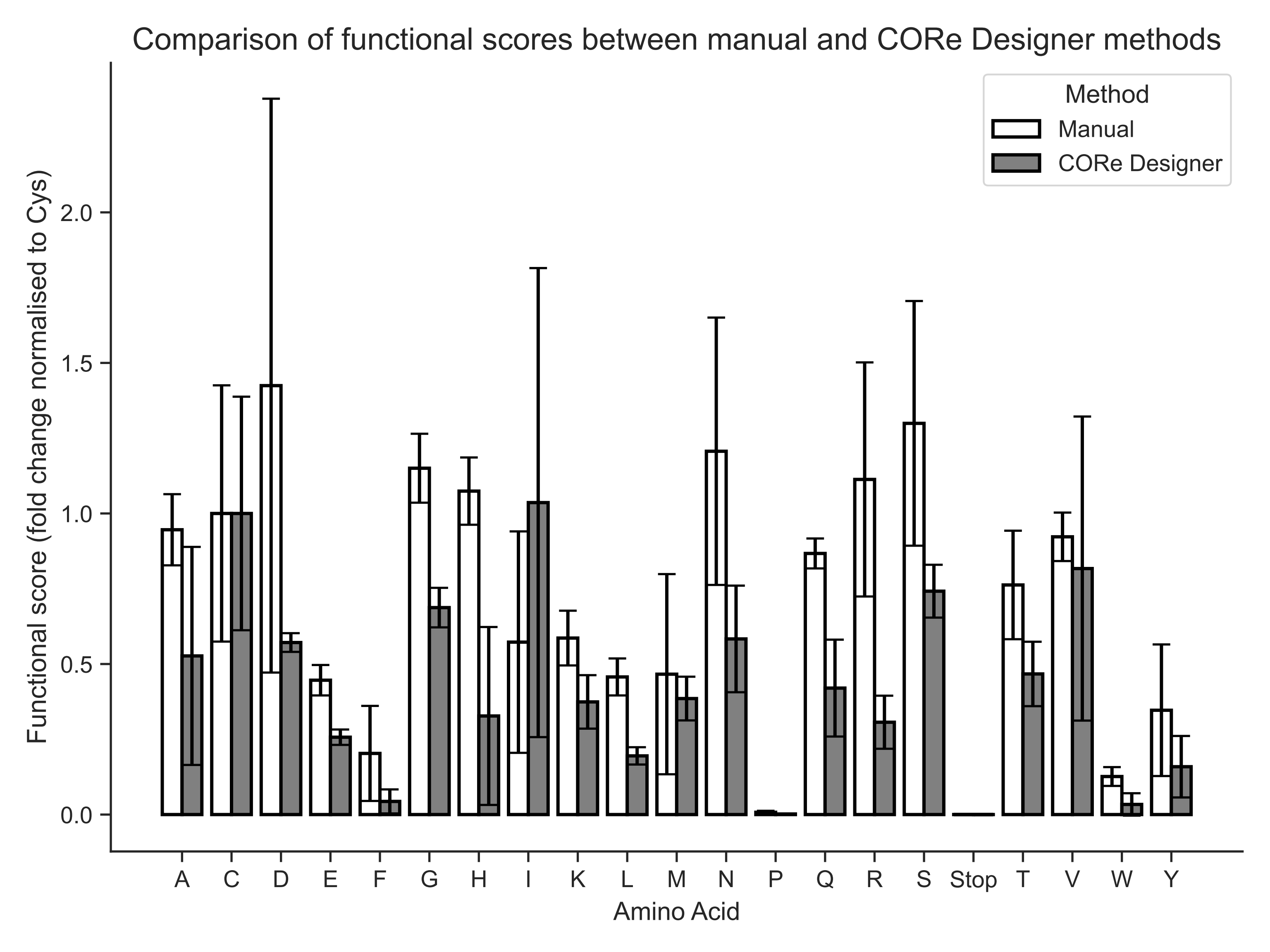

### figure3d.pdf

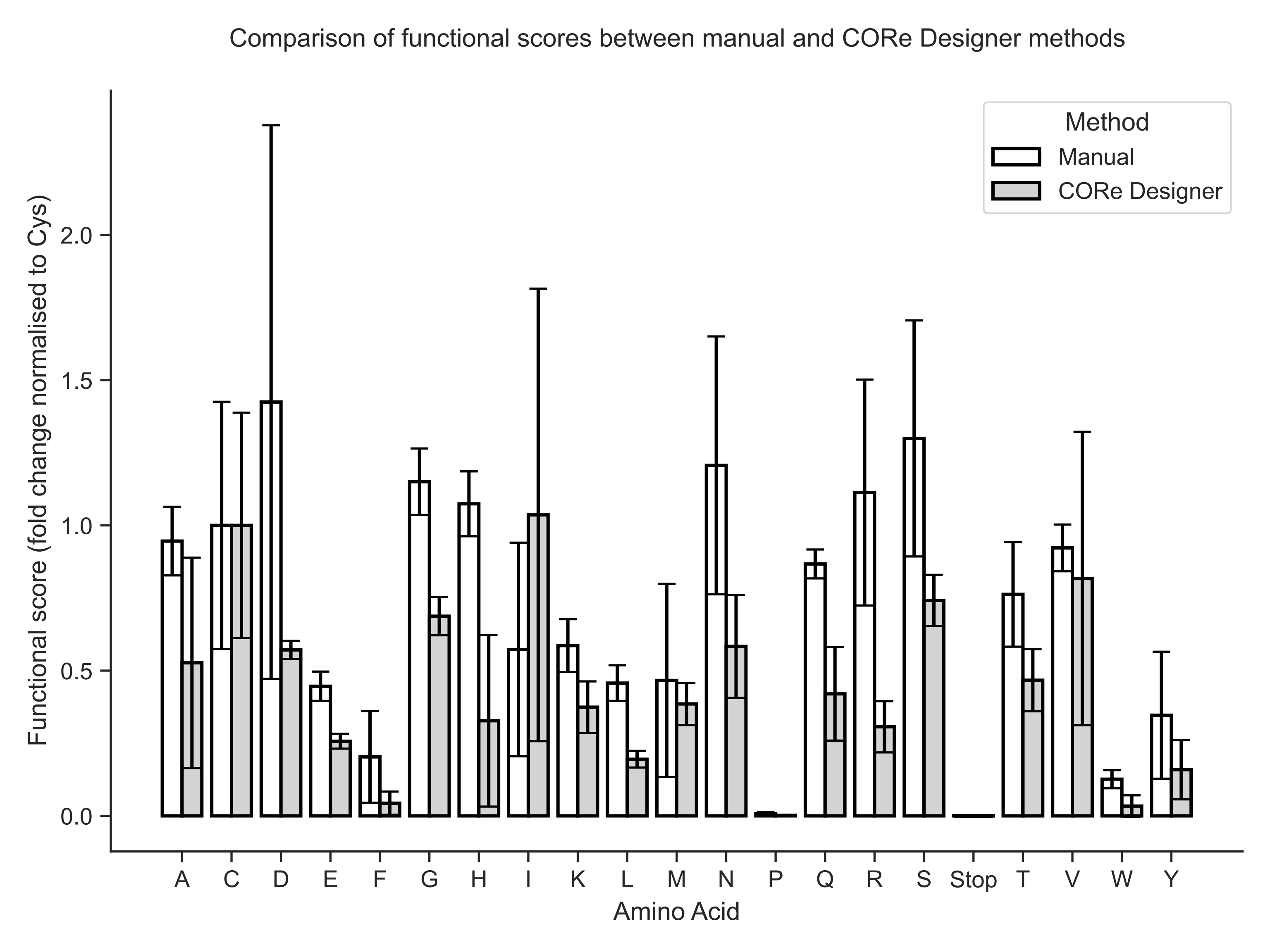

### Figure_2a.pdf

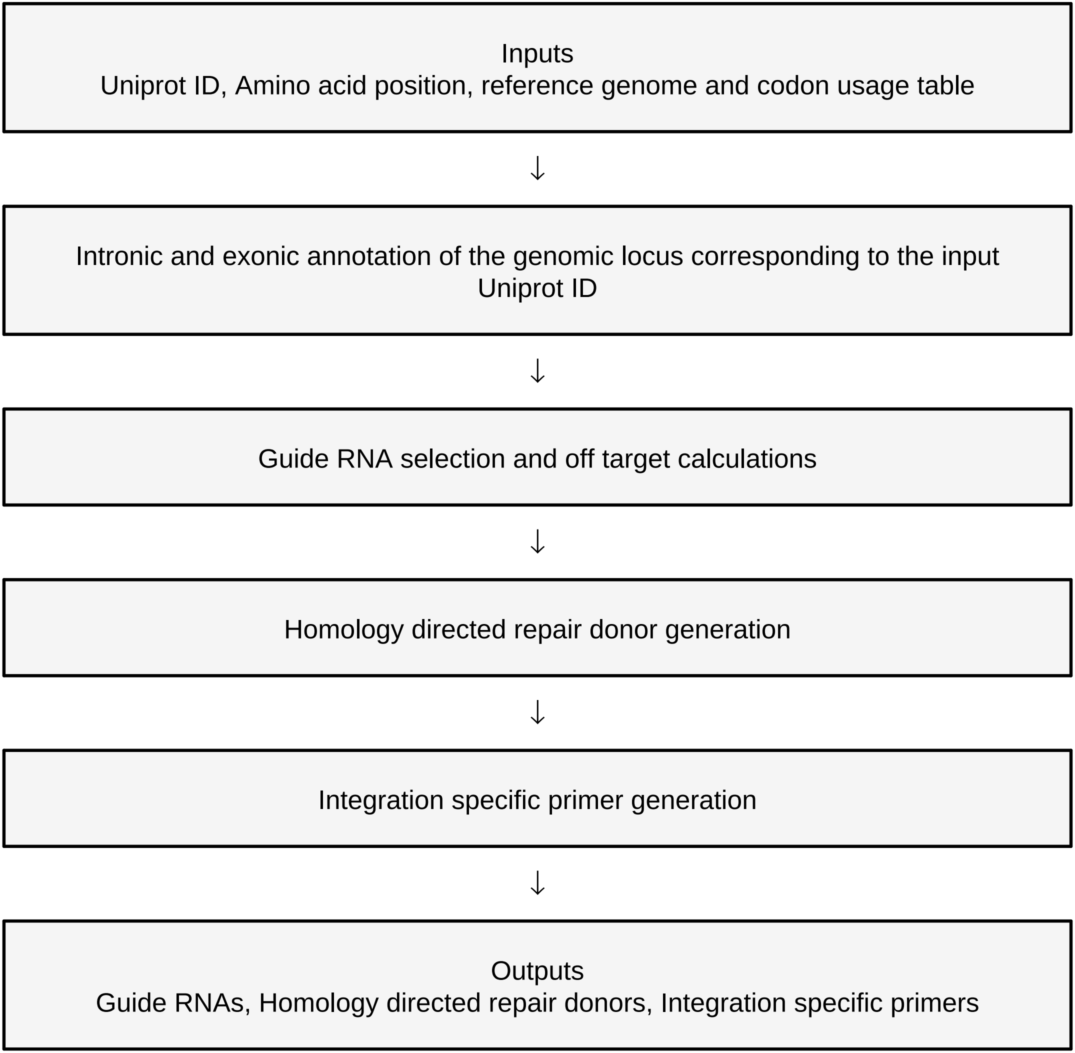

### Figure_2b.pdf

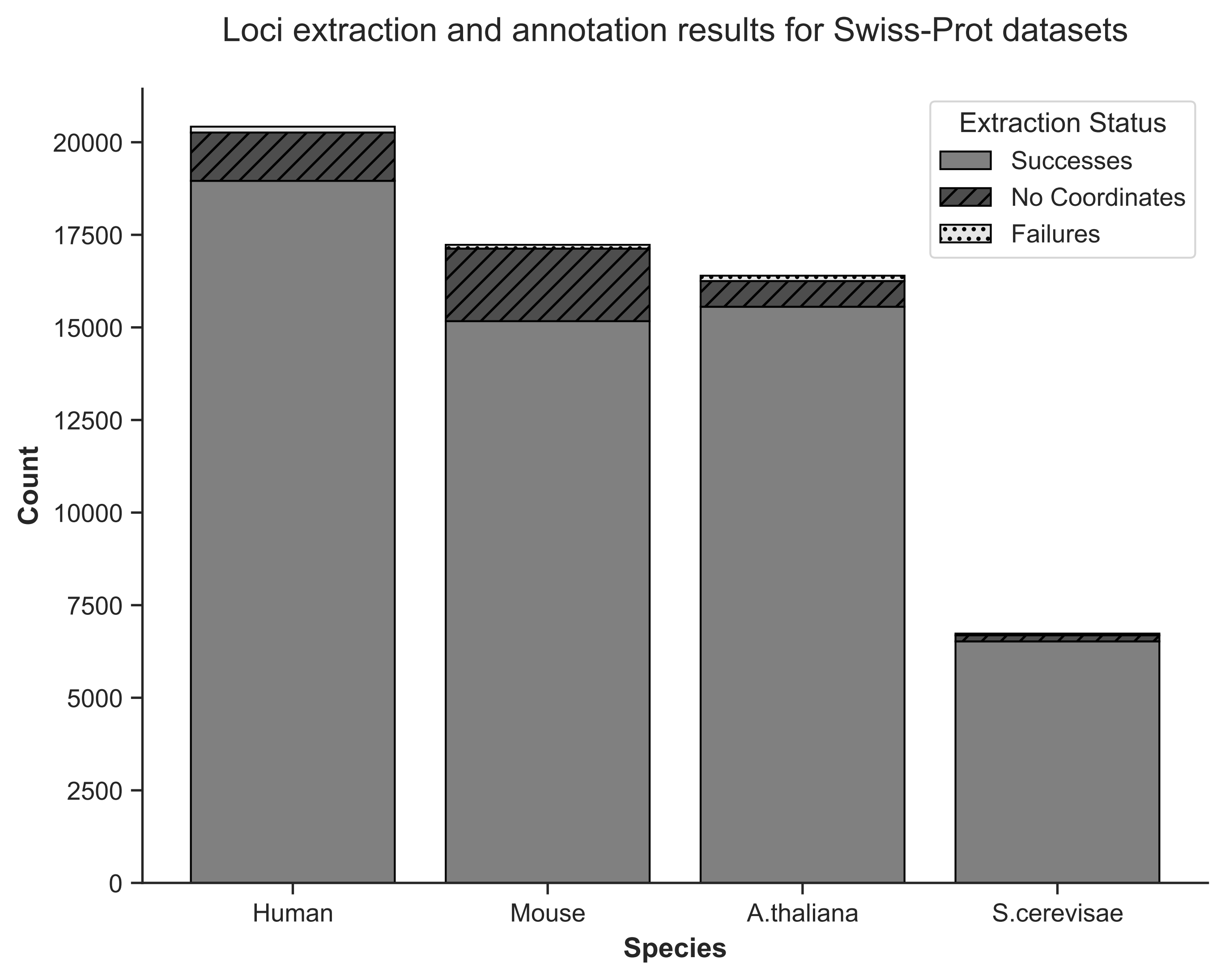

### Figure_2c.pdf

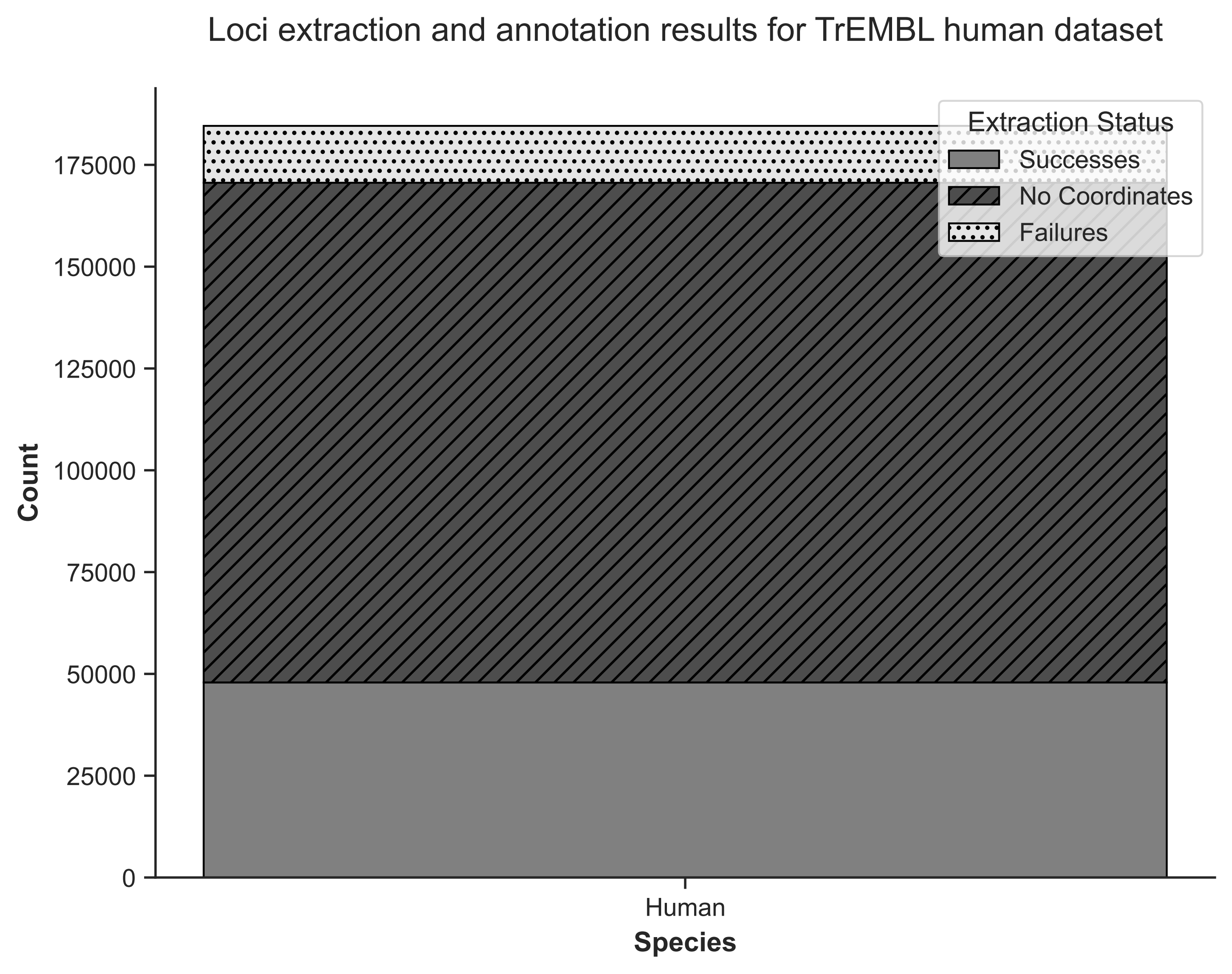

### Figure_2d.pdf

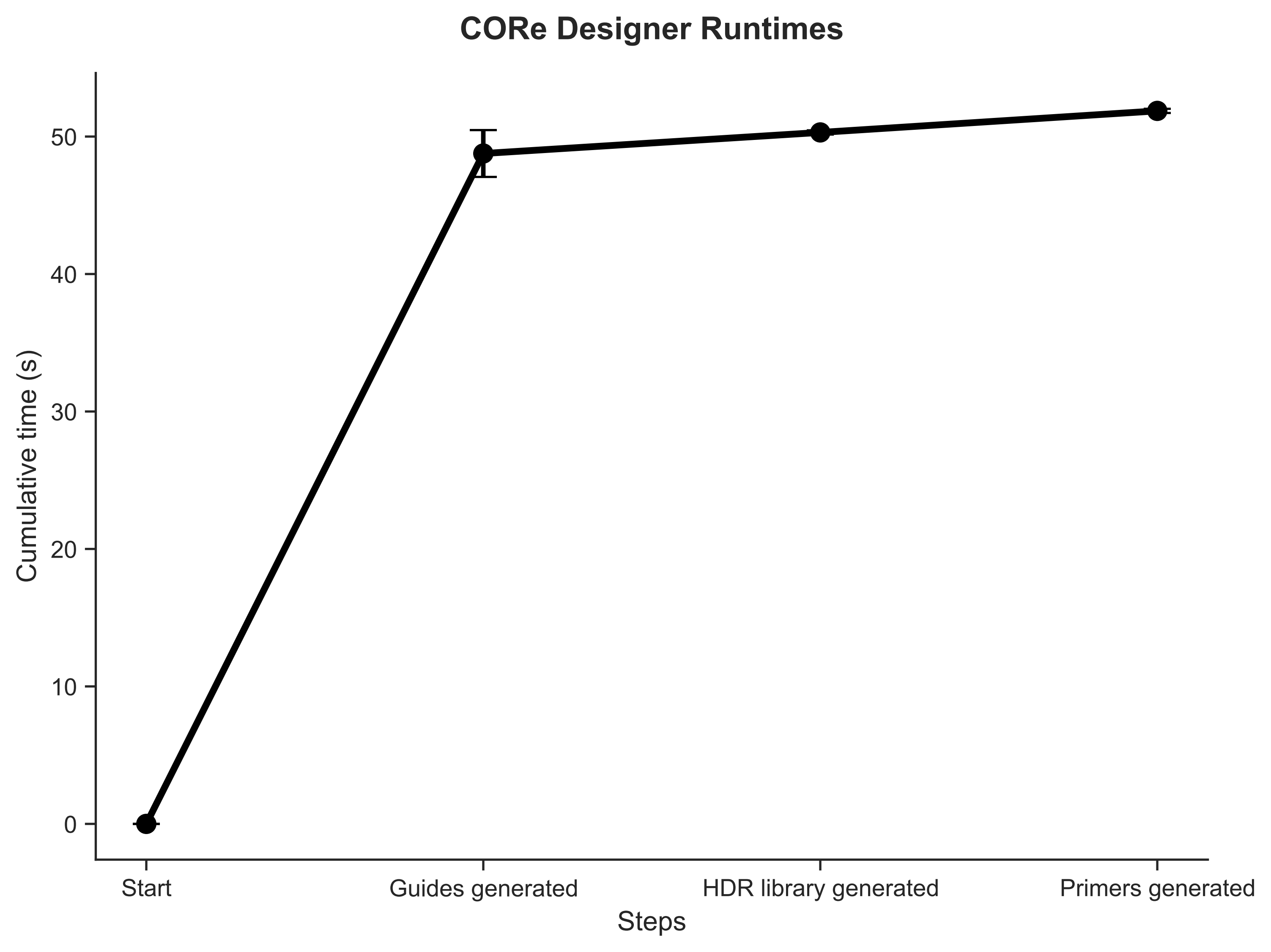

### Figure_2e.pdf

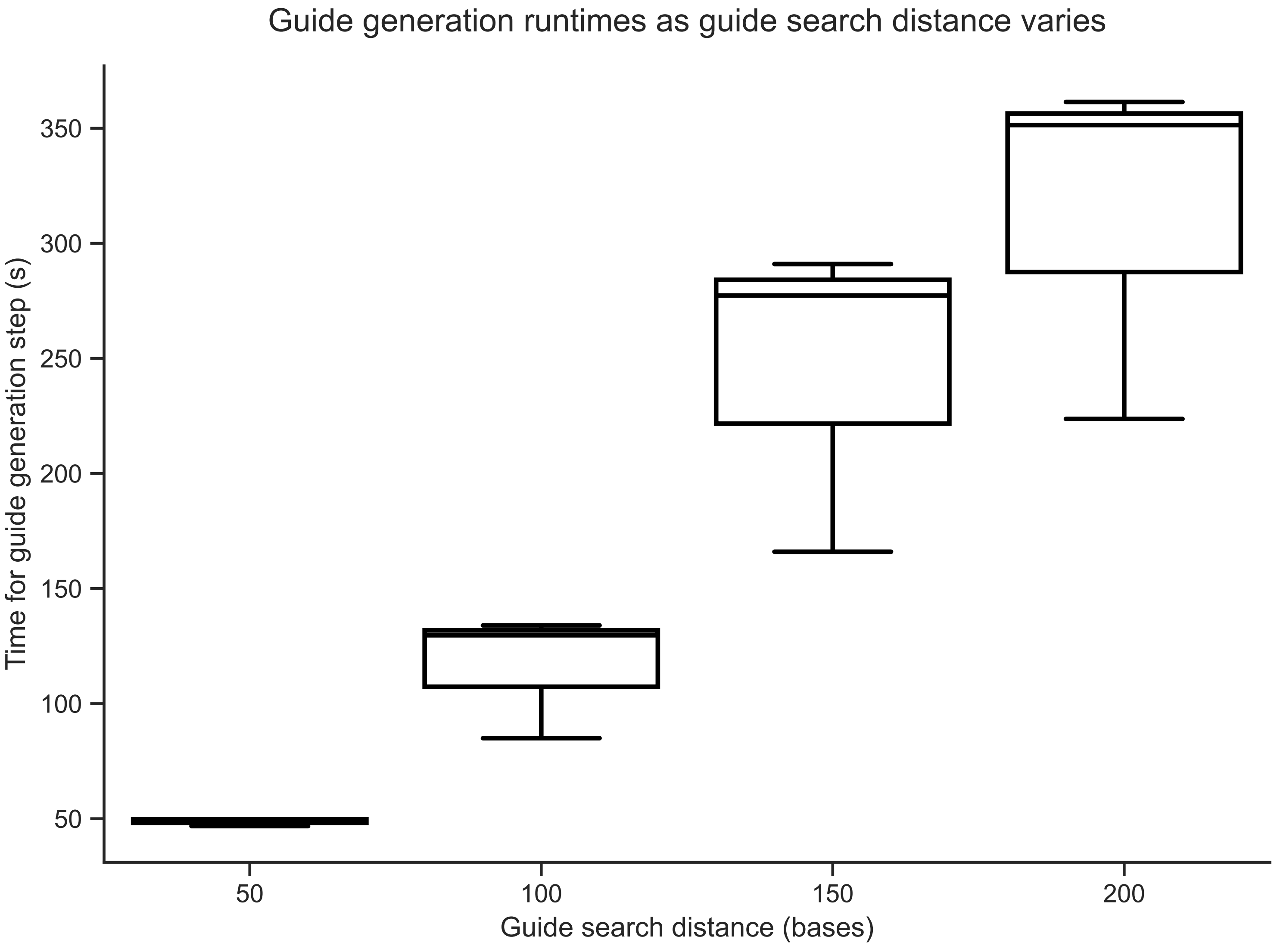

### Figure_3a.pdf

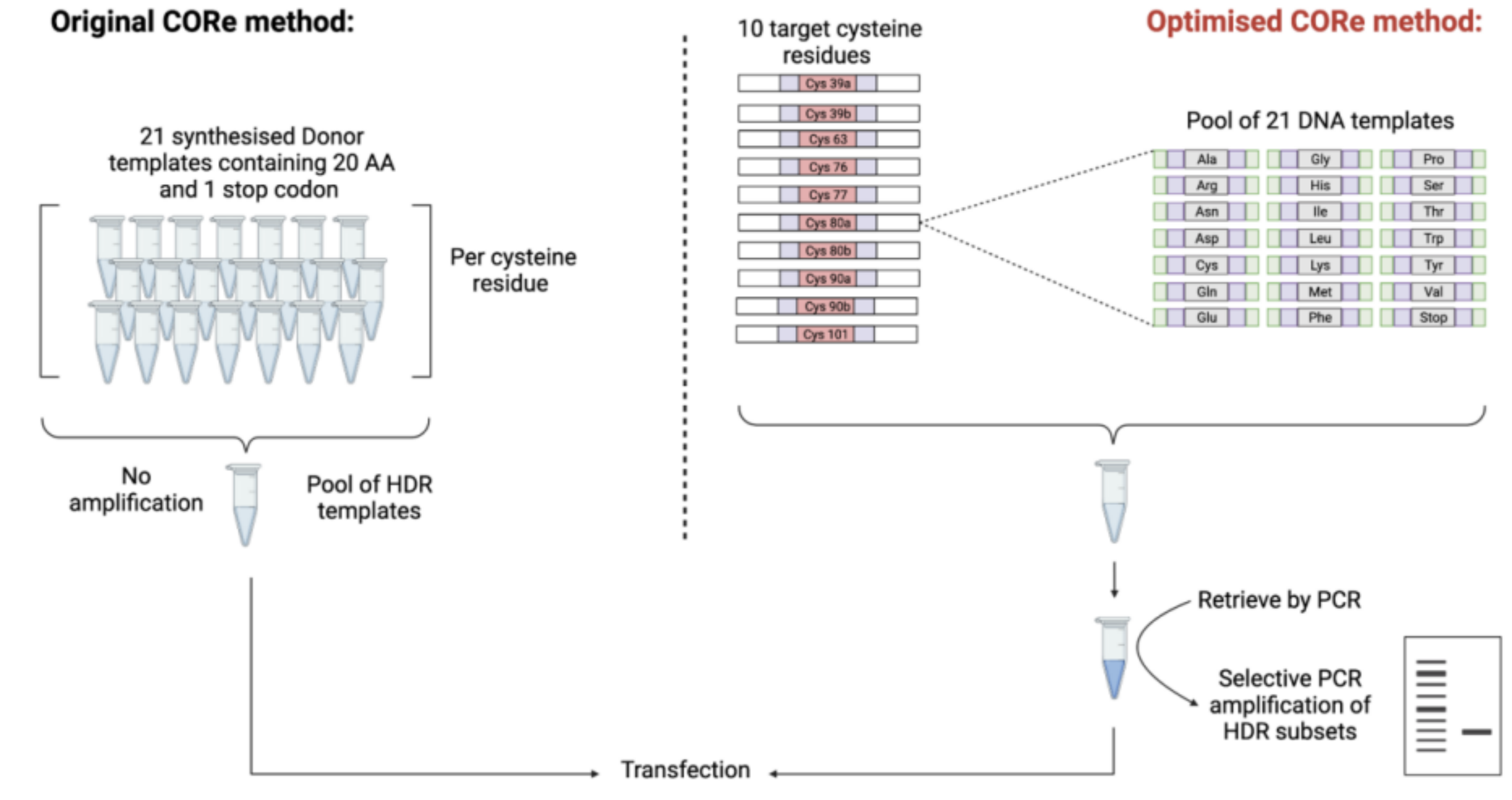

### Figure_3b.pdf

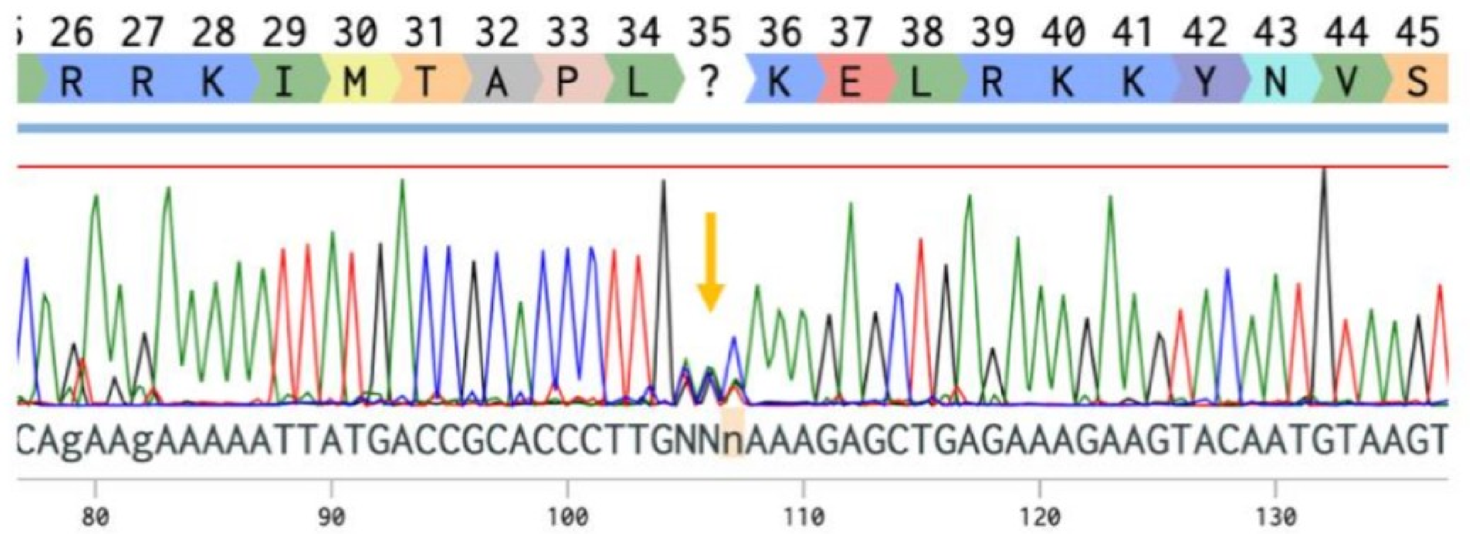

### Figure_3d.pdf

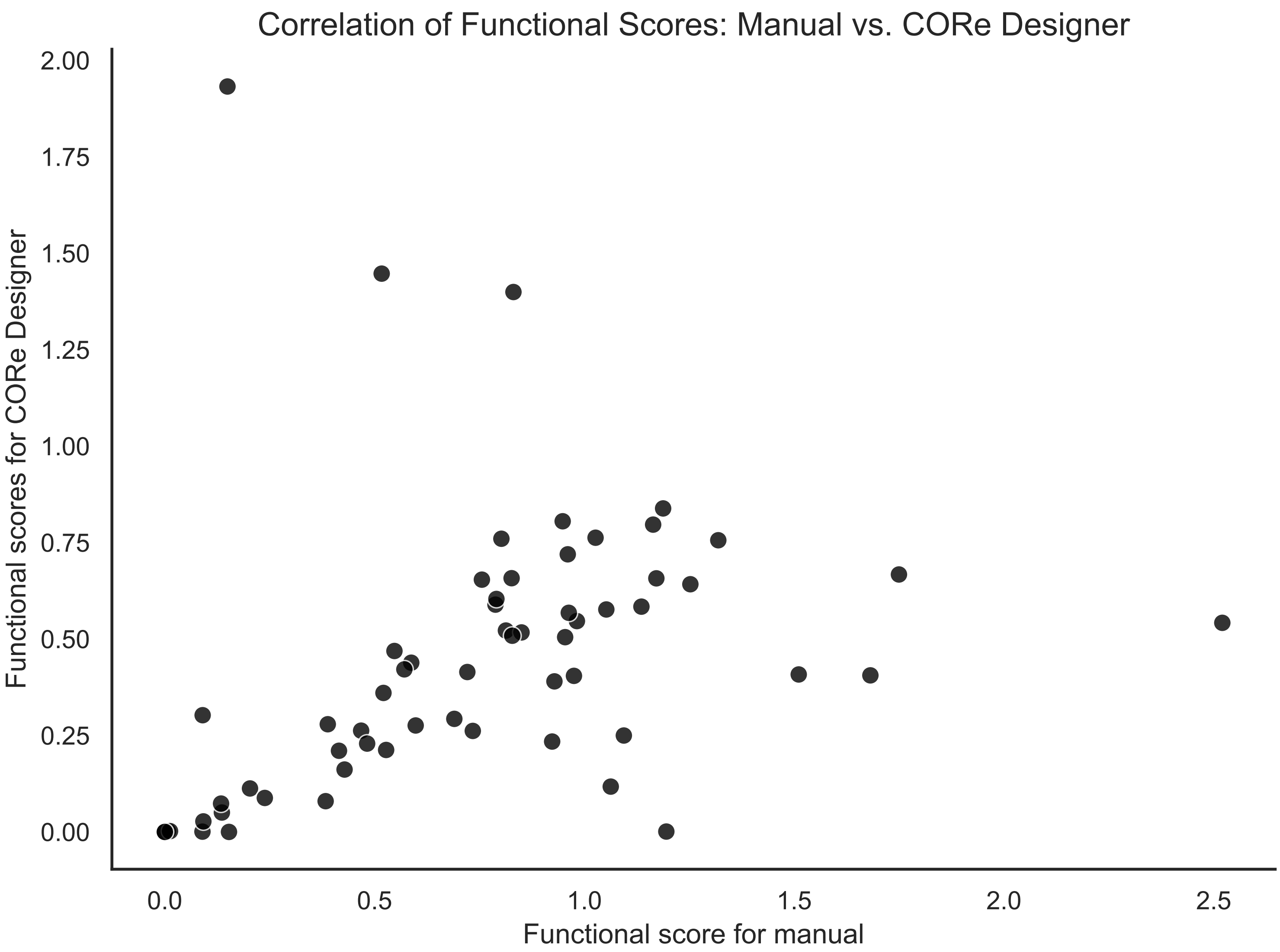
